# Vascular specification from diversified mesoderm culminates in endothelial convergence

**DOI:** 10.64898/2026.10.25.761646

**Authors:** Raz Ben-Yair, Hernan Rubinstein, Yoav Mayshar, Tal Sokolov, Stephanie Gehrs, Liat Fellus-Alyagor, Dana Hirsch, Netta Reines, Ayelet-Hashahar Orenbuch, Saifeng Cheng, Hellmut G. Augustin, Karina Yaniv, Amos Tanay, Yonatan Stelzer

**Affiliations:** Department of Molecular Cell Biology, Weizmann Institute of Science, Rehovot, Israel; Department of Computer Science and Applied Mathematics, Weizmann Institute of Science, Rehovot, Israel; Division of Vascular Oncology and Metastasis, German Cancer Research Center (DKFZ), Heidelberg, Germany; Department of Vascular Biology and Tumor Angiogenesis, European Center for Angioscience (ECAS), Medical Faculty Mannheim, Heidelberg University, Mannheim, Germany; Department of Veterinary Resources, Weizmann Institute of Science; Westlake Laboratory of Life Sciences and Biomedicine, School of Life Sciences, Westlake University, Westlake Institute for Advanced Study, Hangzhou, Zhejiang, China; Department of Immunology and Regenerative Biology, Weizmann Institute of Science, Rehovot, Israel; Helmholtz-Institute for Translational AngioCardioScience (HI-TAC) of the Max Delbrück Center for Molecular Medicine in the Helmholtz Association (MDC) at Heidelberg University, Mannheim, Germany

## Abstract

Vasculogenesis establishes the embryonic circulatory system, yet how a common vascular program emerges across multiple embryonic and extraembryonic developmental environments remains incompletely understood. Here, we reconstruct mouse vasculogenesis through integrated single-cell, spatial, lineage and perturbation analyses. We find that vascular specification emerges from distinct sources of pre-specified mesoderm through a transient Kdr+Etv2− vascular-primed state promoted by BMP signaling. Primed cells persist until E8.0-E8.5, after which they resolve into committed derivatives. Dual blood-endothelial potential is largely restricted to early vitelline progenitors, whereas embryonic and umbilical progenitors contribute predominantly to endothelial lineages. Temporal VEGF signaling perturbations validate the inferred specification sequence, placing VEGF downstream of vascular priming and demonstrating its indispensable role in vascular and primitive blood specification. Unlike most developmental lineages, endothelial derivatives converge toward a common transcriptional state while retaining transcriptional traces of their developmental origins. Together, these findings establish a mechanistically validated developmental sequence for mammalian vasculogenesis.

## INTRODUCTION

In embryonic development, there is a stage at which the embryo’s rising metabolic demands require the establishment of a functional vascular system. In mammalian embryos, vascularization begins in the extraembryonic yolk-sac tissue along with localized formation of blood islands containing primitive blood^1–5^. Later in development, the umbilical system becomes predominant, enabling more efficient exchange through an indirect interface with the maternal circulation^6^. During this period, the embryo begins to develop somatic vasculature. Once the vasculature connects to the heart, initially through the paired dorsal aortae and three venous drainages, blood begins circulating in the system. Interestingly, both blood and endothelial cells (EC) of the vasculature wall emerge from mesoderm through closely related transcriptional programs, previously termed hematoendothelial (HE) state^7,8^. The close developmental relationship between these lineages is further emphasized by the emergence of hematopoietic cells directly from hemogenic endothelium during later embryogenesis^9–13^. However, despite the strong transcriptional and developmental association between blood and endothelium, the existence of an obligate bipotent progenitor generating both lineages *in vivo* remains uncertain^4,14–17^. They may instead originate from progenitors with broader developmental potential or through a transient intermediate population that remains incompletely characterized.

Endothelial and vascular diversity across tissues and organs is now increasingly characterized by single-cell transcriptomic atlases defining organotypic endothelial identities^18–23^, arterial-venous specialization^24^, and mature vascular heterogeneity^4,8,18–23,25–29^. In contrast, vasculogenesis, the *de novo* emergence of endothelial lineages from mesoderm during gastrulation, remains poorly understood within the continuous and parallel diversification of mesoderm across embryonic and extraembryonic compartments^30–42^. We have previously established a temporally resolved single-cell model of mouse gastrulation, enabling reconstruction of continuous and parallel differentiation trajectories across embryonic and extraembryonic tissues^30,43^. Interestingly, using a computational optimal transport approach, the model inferred multiple mesodermal origins converging to a common hematoendothelial progenitor state.

Inspired by this intriguing computational prediction, we here characterize the spatiotemporal sequence of endothelial specification from distinct mesodermal trajectories during gastrulation. BMP and VEGF signaling act early within these trajectories, preceding overt endothelial differentiation and progressively restricting mesodermal potential toward endothelial fate. Lineage analyses identify a developmental transition after Embryonic day (E) 8.0, separating plastic cell states capable of generating both endothelial and alternative mesodermal derivatives from more committed vascular progenitors. Despite their distinct developmental origins, these trajectories ultimately converge toward a near-uniform endothelial transcriptional state.

## RESULTS

### Construction of a spatiotemporally informed vascular atlas of gastrulation

To begin characterizing the developmental sequence of endothelial specification, we defined an HE transcriptional module comprising genes characteristic of both endothelial and blood progenitors^7,30,44^, including *Kdr*, *Etv2*, *Gata2*, *Tal1,* and *Fli1* (**Figure 1A**). Quantification of this module across the gastrulation atlas revealed that, whereas endodermal and ectodermal populations, which do not contribute to the vascular lineage, consistently exhibited minimal HE-associated gene expression, both embryonic and extraembryonic mesoderm express these genes variably (**Figure 1A,B**). These observations suggested that subsets of diversified mesoderm populations may reside along developmental trajectories leading toward endothelial and hematopoietic fates. Notably, caudal mesoderm, which has developmental origins distinct from early primitive streak-derived mesoderm, consistently exhibited low HE module activity, suggesting it is not part of the vascular trajectory during gastrulation. Consistent with these observations, analysis of our previously established time-resolved transcriptional flow model of mouse gastrulation^30^ predicts that HE states emerge from multiple mesodermal trajectories that have already diversified (**Figure 1C**). These included early contributions from the earliest mesoderm to exit the primitive streak (nascent mesoderm), followed by extraembryonic, yolk-sac, cardiopharyngeal, rostral, and paraxial mesoderm trajectories.

**Figure 1.**
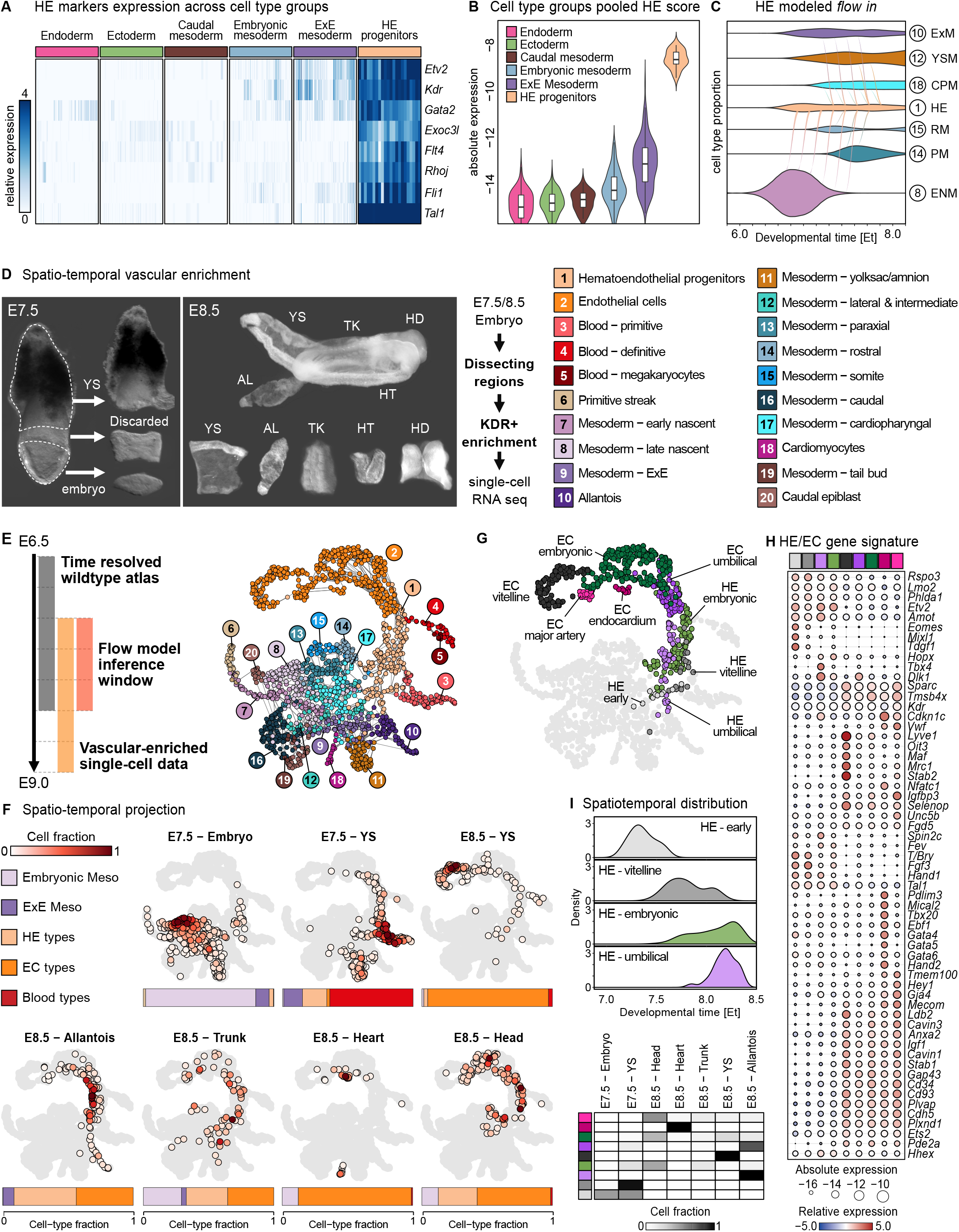
Construction of a spatio-temporally informed, vascular-enriched atlas. **A,** Heatmap of HE-module marker gene expression across cell type groups (endoderm, ectoderm, caudal mesoderm, embryonic mesoderm, ExE mesoderm and HE progenitors). Color indicates relative expression. **B,** Violin plots of the HE-module score (Log_2_). Embedded box plots show the median and inter-quartile range. **C,** Transcriptional flow model predicting the relative contribution of distinct mesodermal subtypes to HE progenitors over developmental time (Et). Colored by cell type (numbered key). **D,** Representative E7.5 and E8.5 embryos dissected into spatially defined compartments, enriched for KDR-positive cells and profiled by scRNA-seq (workflow, right). **E,** Sources of the integrated single-cell data (WT temporal atlas, and vascular-enriched data; left) and the resulting vascular-enriched manifold (right), colored by cell type (numbered key), each dot represents a MetaCell. **F,** Manifold enrichment representation of Kdr sorted cells per dissected compartment (as in d). MetaCell color represents relative contribution of cell types. Bars below each projection show the relative abundance of the main cell types per compartment. **G,** Atlas projection of the re-annotated vascular subtypes. HE and EC types are color-coded, non-vascular types are grey. **H,** Dot plot of marker genes defining the HE and EC subtypes; dot size indicates absolute expression (Log_2_); color, relative expression. **I,** Spatiotemporal distribution of the HE subtypes. Density plots showing developmental-time distribution of HE subtypes (top) and a heatmap representing vascular subtype proportions across spatial compartments (bottom). HE, hematoendothelial progenitors; EC, endothelial cell; ENM, early nascent mesoderm; ExM, extraembryonic mesoderm; YSM, yolk-sac mesoderm; CPM, cardiopharyngeal mesoderm; RM, rostral mesoderm; PM, paraxial mesoderm; YS, yolk sac; AL, allantois; TK, trunk; HT, heart; HD, head.

To gain further insight into this process, we constructed a vascular-enriched single-cell atlas of gastrulation. E7.5 and E8.5 embryo were dissected into spatially segregated embryonic, yolk-sac, allantois, trunk, heart and head compartments, dissociated into single-cell suspensions, immunolabeled using a KDR/VEGFR2 antibody, and enriched by FACS sorting for vascular-associated populations prior to scRNA-seq profiling using MARSeq (**Methods**, **Figure 1D**). We then integrated these data with all mesodermal and extraembryonic mesodermal populations from our wild-type (WT) gastrulation atlas, generating a combined vascular-enriched manifold consisting of 46,265 cells grouped into 1,656 MetaCells (MCs), providing robust statistical representation of continuous developmental states. The resulting atlas spans primitive streak and mesodermal populations through HE states and differentiated endothelial and hematopoietic lineages (**Figure 1E**).

### Kdr expression marks broad mesodermal and vascular populations

Interestingly, at E7.5, most KDR-positive cells derived from the embryo-proper compartment (94.5%) mapped to broad mesodermal populations rather than HE or endothelial states (**Figure 1F**). Thus, initial KDR expression marks broad mesodermal populations before commitment toward vascular fates. In contrast, KDR-positive cells isolated from the yolk-sac at E7.5 were enriched for primitive blood, HE and extraembryonic mesodermal states (ExM) (64%, 18.5%, 15.2%, respectively). By E8.5, KDR-positive cells predominantly mapped to HE (23%), and endothelial populations (66.4%) as well as definitive blood (1.3%), whereas mesodermal representation became substantially reduced (9.3%). Using integrated spatial, temporal and gene expression profiles, we annotated four HE subtypes (early, vitelline, embryonic and umbilical), five EC subtypes (vitelline, embryonic, umbilical, major artery and endocardium) and three blood subtypes (primitive, definitive and megakaryocytes) (**Figure 1G-H** and **S1A-B**).

The earliest HE cells, predominant between E7.0 and E7.5 in both the embryonic compartment and the yolk-sac were characterized by persistent expression of early nascent mesoderm genes such as *T*, *Mixl1*, *Eomes*, and *Tdgf1*. Vitelline HEs emerged after E7.5 in the yolk-sac, characterized by extraembryonic genes such as *Fgf3* and *Hand1* and expression of highly specific transcription factors (TFs) such as *Runx1*. Umbilical HEs appeared later (after E8.0) and express an allantois program, including *Tbx4* and *Dlk1*. Embryonic HEs were defined primarily by their spatial location. They lacked a unique, defining transcriptional signature and represent a heterogeneous population (see below) with broad temporal distribution (**Figure 1H-I**).

Vitelline ECs expressed a unique signature including genes like *Lyve1*, *Oit3*, *Maf*, *Mrc1,* and *Stab2* (**Figure 1H**). Consistent with this annotation, immunofluorescence analysis demonstrated spatially restricted LYVE1 expression within vitelline endothelial structures (**Figure S1B**). Umbilical ECs maintained significant levels of *Dlk1*, while endocardial ECs expressed a unique cardiac gene program including *Gata4,5,6*, *Hand2*, and *Tbx20*. Embryonic ECs, similar to their HE counterparts, were heterogeneous and defined based on spatial context. Major artery ECs were predominantly located in the head compartment at E8.5, consistent with the initially anterior localization of the dorsal aorta, and expressed characteristic genes such as *Unc5*, *Hey1*, *Gja4,5,* and *Notch4*. Finally, primitive and definitive blood populations were distinguished by distinct transcriptional programs. Primitive blood expressed genes including *Cd24a, Cdh23,* and residual *Fgf3*, whereas definitive blood was characterized by expression of *Myb, Cd47*, *Cd34* and the embryonic hemoglobin gene *Hbb-y*. Megakaryocyte progenitors were identified by expression of platelet-lineage genes including *Mpl*, *Pf4, Itgb3/Cd61,* and *Thbs1*. Blood and endothelial populations were further distinguished by divergent regulation of HE genes. In both lineages, *Etv2* expression was strongly repressed following specification. However, whereas *Tal1* expression was maintained and further increased in blood lineages, it progressively declined during endothelial differentiation (**Figure 1H** and **S1C-D**).

### Hematoendothelial specification follows diversification of mesodermal identities

The vascular-enriched atlas described above integrates cells collected across multiple embryos, developmental stages and anatomical compartments, precluding direct reconstruction of developmental flows. To reconstruct developmental trajectories leading to individual HE populations, we projected our previously established time-resolved WT gastrulation model^30^ onto the vascular-enriched atlas and generated a higher-resolution flow representation of vascular specification (**Methods**). Flow analysis linked distinct HE populations with different mesodermal trajectories (**Figure 2A-C**). Vitelline HEs were linked to early HEs, which in turn were associated with nascent mesoderm. Umbilical HEs are mapped to the allantois trajectory, itself originating from extraembryonic mesoderm. Embryonic HEs were linked to multiple embryonic mesoderm populations, including rostral, cardiopharyngeal and paraxial mesoderm.

**Figure 2.**
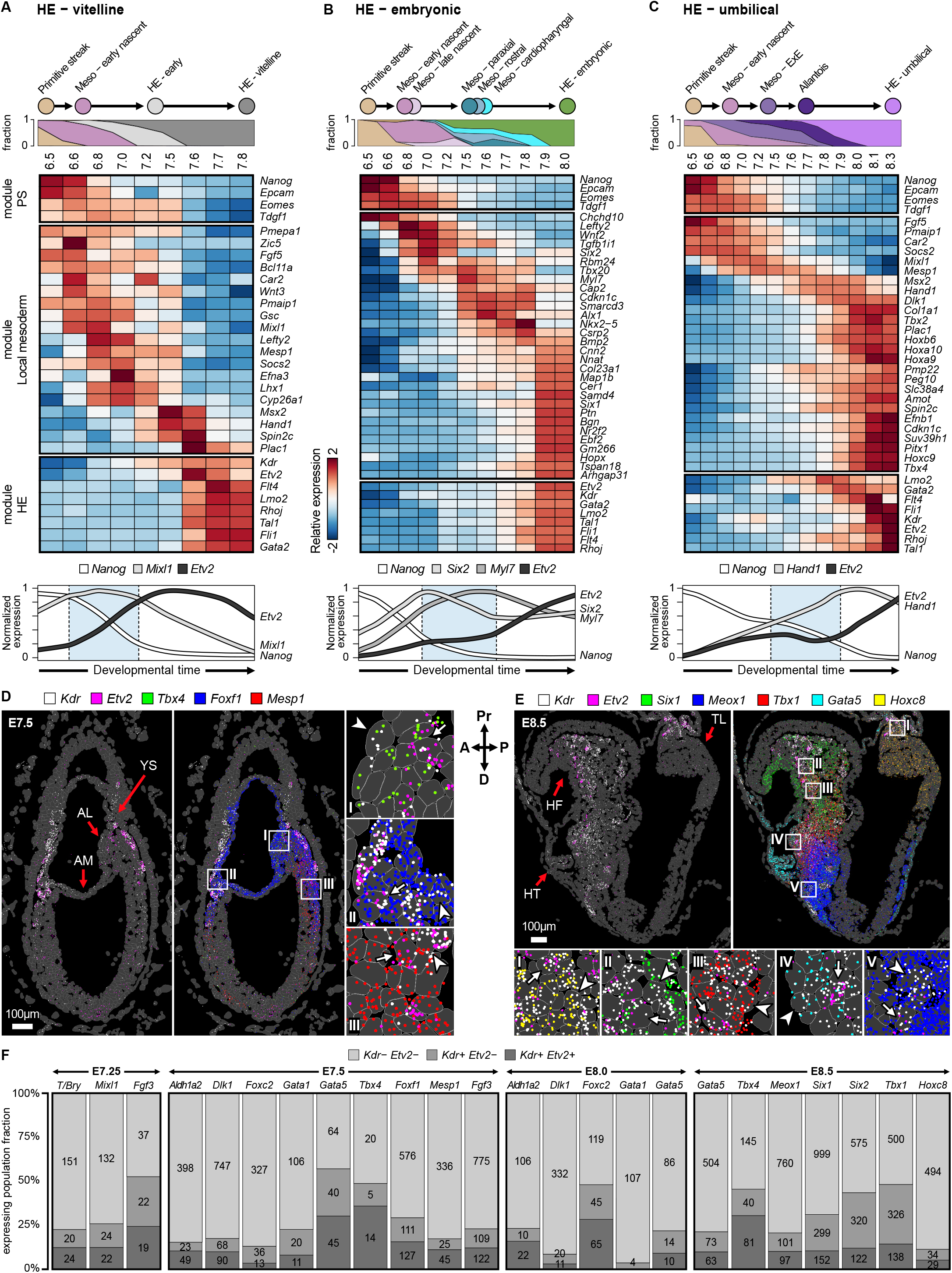
Cell state dynamics in vascular differentiation. **a-c,** Transcriptional dynamics along the three main vascular trajectories: vitelline (**A**), embryonic (**B**) and umbilical (**C**) HE. Top, trajectory schematic and cell type proportions over developmental time (Et). Heatmaps show the relative expression of stage-defining genes grouped into primitive-streak, local-mesoderm and HE modules (left). Bottom, normalized expression of *Etv2* versus representative primitive-streak and local-mesoderm markers over time. Mesoderm-to-HE transitional window marked with a blue box. **D,** MERSCOPE section of an E7.5 embryo showing *Kdr* (white) and *Etv2* (magenta), together with the local-mesoderm markers *Tbx4* (green, allantois), *Foxf1* (blue, ExM), and *Mesp1* (red, nascent mesoderm), at single-cell resolution. Markers are detected alone (local mesoderm), co-expressed with *Kdr* (vascular-primed mesoderm, white arrowhead), or with *Kdr* and *Etv2* (specified HE, white arrow). Boxed insets are magnified on the right. **E,** E8.5 embryo section showing the distribution of local-mesoderm markers *Six1* (green, rostral mesoderm), *Meox1* (blue, paraxial mesoderm), *Tbx1* (red, cardiopharyngeal mesoderm), Gata5 (cyan, cardiopharyngeal mesoderm), and Hoxc8 (yellow, caudal mesoderm). **F,** Stacked-bar quantification of MERSCOPE data across four stages (E7.25, E7.5, E8.0 and E8.5), showing, for each local-mesoderm marker, proportions of expressing cells that are *Kdr*-/*Etv2-* (local mesoderm), *Kdr*+ /*Etv2*- (vascular-primed mesoderm) or *Kdr*+/*Etv2*+ (specified HE). Numbers indicate cell counts. A, anterior; P, posterior; Pr, proximal; D, distal; YS, yolk sac; AL, allantois; AM, amnion; HF, head fold; HT, heart; TL, tail.

Gene expression dynamics along these trajectories revealed a stepwise sequence of specification events. Pluripotency and primitive streak genes, including *Nanog* and *Eomes*, were progressively downregulated, followed by activation of transcriptional programs characteristic of locally specified mesoderm. These included extraembryonic and allantoic markers such as *Hand1*, *Msx2,* and *Dlk1* in the umbilical trajectory, and rostral and cardiopharyngeal markers including *Six2*, *Myl7*, *Tbx20*, *Nkx2-5,* and *Smarcd3* in embryonic trajectories. Only subsequently was the HE program activated, marked by induction of *Etv2* together with additional HE markers such as *Lmo2*, *Tal1,* and *Fli1* (**Figure 2A-C**). A potential exception to this pattern is represented by the early HEs, which are predicted to arise directly from migratory nascent mesoderm and therefore lack strong tissue-specific signatures, instead expressing early mesodermal TFs such as *Mixl1* and *Mesp1*.

Analysis of HE- module dynamics further revealed temporal differences between trajectories. In the vitelline trajectory, HE module expression peaked early during differentiation. In contrast, embryonic and allantois trajectories maintained intermediate HE module expression over time before undergoing a sharper induction and transition into HE states, typically after E7.5 (**Figure 2A-C** lower panels and **S2**).

### Spatiotemporal profiling supports HE specification from pre-specified mesoderm

To place the newly defined HE populations in spatial context and examine their relationship with local mesodermal identities, we profiled gastrulating embryos from E7.25 to E8.5 using the MERSCOPE platform (Vizgen), enabling simultaneous single-molecule-level detection of approximately 500 transcripts in tissue sections with cellular resolution (**Methods**, **Figure 2D, E** and **S3**). This analysis revealed three recurrent spatial configurations of mesodermal and HE-associated gene expression. First, broad mesodermal domains expressing local transcriptional programs without detectable *Kdr* or *Etv2*. Second, subsets of these domains expressed *Kdr* while lacking *Etv2*, suggesting an intermediate state in which locally specified mesoderm is associated with vascular competence but has not yet activated the HE program. Third, populations in which *Kdr* and *Etv2* are co-expressed, consistent with HE specification (**Figure 2F**).

This configuration was observed across multiple anatomical contexts. At late streak stage (∼E7.25), *Kdr*-positive cells occupied uniform mesodermal domains markers by *Fgf3, Mixl1* and *T/ Brachyury*, likely corresponding to the early HE population identified in the vascular atlas. At E7.5, *Kdr*-positive cells appeared within mesodermal territories marked by nascent mesoderm (including *Aldh1a2, Foxc2,* and *Mesp1*), yolk-sac mesoderm (including *Foxf1, Dlk1* and *Fgf3*) and nascent allantois (*Dlk1*, *Tbx4*) (**Figure 2D** and **S3A-C**). *Gata1*-positive blood islands-associated cells also expressed considerable levels of *Kdr* at this stage (**Figure 2F** and **S3C**). At E8.0, vascular-associated populations were embedded within increasingly specialized mesodermal domains, including yolk-sac (*Dlk1*), rostral mesoderm (*Foxc2*), cardiopharyngeal mesoderm (*Gata5*) and paraxial mesoderm (*Aldh1a2*) (**Figure S3D**). A distinct organization was observed within the blood islands, where mostly *Etv2*-negative and *Gata1*-positive primitive blood cells gradually decrease *Kdr* co-expression by E8.0 (**Figure 2F** and **S3D**).

Between E7.5 to E8.0, *Kdr/Etv2*-positive population increased in both *Aldh1a2-* and *Foxc2-*positive population, suggesting *de novo* embryonic HE specification (**Figure 2F**). By E8.5, *Kdr*/*Etv2*-positive populations were detected within multiple spatially distinct domains corresponding to committed mesodermal populations (**Figure 2E** and **S3E-F**). These included cardiac mesoderm (*Gata5*), head mesenchyme (*Six1*, *Six2,* and *Tbx1*), paraxial mesoderm (*Meox1*), the developing allantois (*Tbx4*), and a distinct caudal population expressing *Hoxc8*. Thus, HE specification was observed in the context of already established regional mesodermal identities rather than preceding them. A potential exception is represented by the early HEs, which are predicted to arise directly from migratory nascent mesoderm and therefore lack strong tissue-specific signatures. Consistent with this interpretation, proximal embryonic regions at E7.25-7.5 exhibited early vascular signatures that were spatially continuous with yolk-sac HEs (**Figure 2D, F** and **S3A-C**).

The diversity of spatial contexts associated with embryonic HEs revealed substantial heterogeneity within this population, consistent with their heterogeneous transcriptional profile and compound flow prediction (**Figure 1H** and **Figure 2B**). To characterize this population further, we returned to the vascular atlas, which provides greater transcriptional depth and temporal resolution. Embryonic HEs could be subdivided into early, caudal and rostral populations exhibiting distinct transcriptional signatures and spatiotemporal profiles (**Figure S4A**). Early embryonic HEs retained expression of nascent mesoderm genes such as *T* and *Mixl1* (**Figure S4B**), whereas caudal HEs expressed genes resembling allantoic mesoderm, including *Hoxc8* and *Dlk1*. Rostral HE populations expressed markers of rostral and cardiopharyngeal mesoderm such as *Tbx1* and *Col2a1*, while more mature rostral populations exhibited increasing expression of endothelial regulators including *Pecam1*, *Erg*, *Icam2* and *Nos3*^20^. Together, transcriptional flow analysis and spatiotemporal profiling support a model in which HE specification emerges independently within multiple pre-specified mesodermal populations distributed across embryonic and extraembryonic compartments, following local mesoderm diversification.

### Endothelial specification culminates in extensive but incomplete transcriptional convergence

We next asked how the distinct mesodermal origins of HE populations are reflected in differentiated endothelial states. Comparison of transcriptional similarity across vascular populations revealed marked differences between HE and EC states (**Figure 3A**). Whereas HE populations exhibited substantial transcriptional divergence, EC populations clustered tightly despite originating from multiple HE subtypes, indicating extensive transcriptional convergence during endothelial differentiation. To characterize the basis of this convergence, we analyzed the time and extent of transcriptional change characterizing the transition between local mesoderm types and their corresponding HE and EC derivatives (**Figure 3B** and **S5A**). Mesodermal programs progressively declined following HE specification, whereas endothelial programs were activated across all trajectories. These dynamics were particularly pronounced in the early differentiating vitelline lineage, where nascent mesoderm signatures diminished significantly in the transition between early to vitelline HEs. TFs such as *Mixl1*, *Meox1,* and *Msx1*, which characterize the mesodermal progenitors of vitelline, embryonic subsets, and umbilical lineages respectively, showed progressive loss of expression during differentiation through HE states, culminating in complete repression in ECs. Despite this overall trend, a subset of genes exhibited sustained expression, representing transcriptional “memory” that persists in ECs. For instance, *Foxf1*, *Foxc2,* and *Dlk1* persist throughout the vitelline, embryonic, and umbilical lineages, respectively (**Figure 3C**).

**Figure 3.**
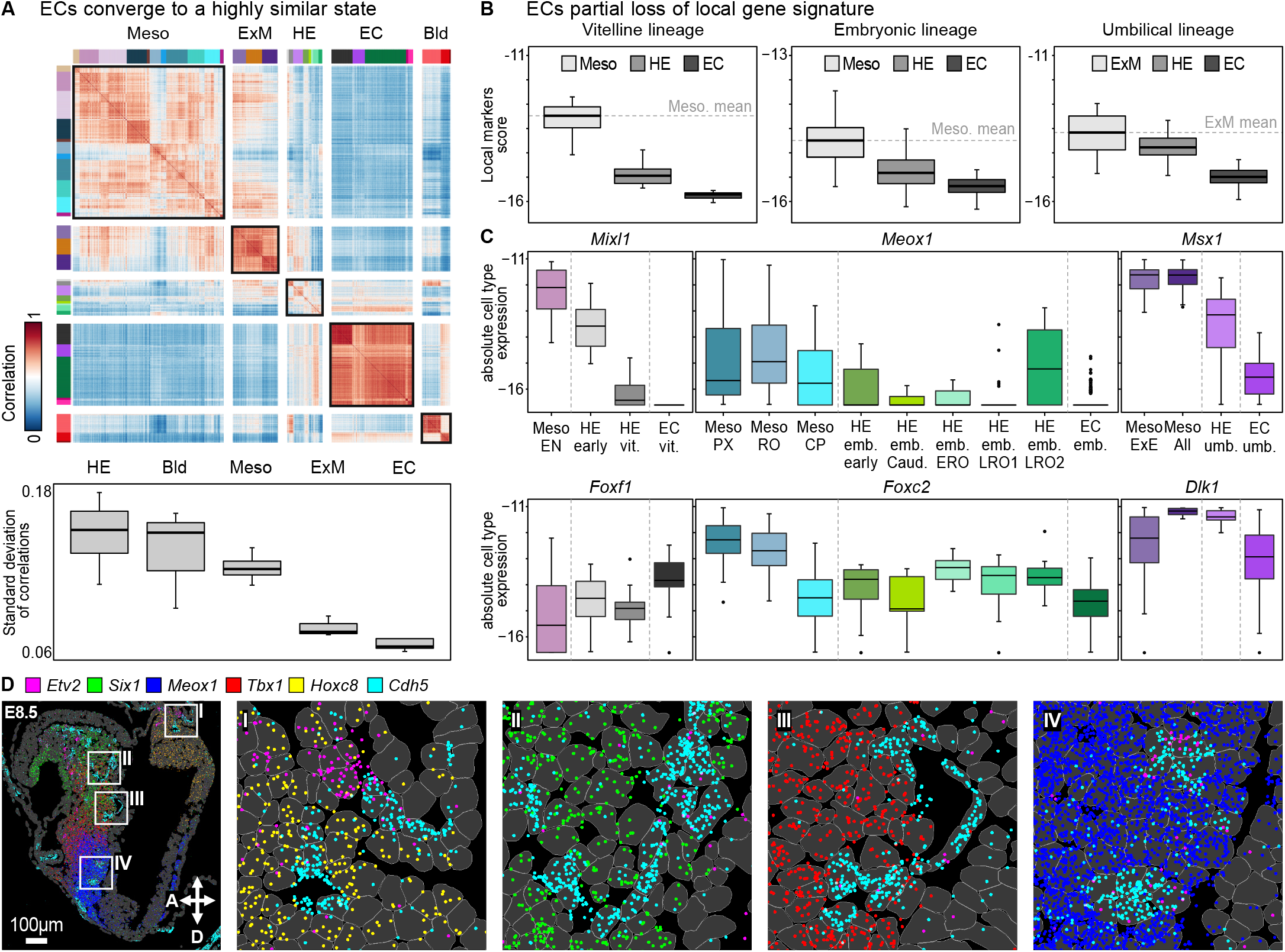
Incomplete loss of local mesoderm signature in ECs during vascular differentiation. **A,** Correlation matrix of transcriptional similarity within major cell type groups in the atlas. Bottom, box plots quantifying the overall variation (standard deviation of correlations) within the major cell type groups. **B,** Box plots of local-marker score (Figure S5A) showing gradual loss of typical local-mesoderm markers along the three main vascular lineages (vitelline, embryonic and umbilical), comparing mesoderm, HE and EC; grey dashed line represents mesoderm (or ExM) mean score. **C,** Representative absolute expression box plots of key local-mesoderm markers across the three vascular lineages (color-coded), showing gradual loss (top) or conservation (bottom). **D,** MERSCOPE section of a representative E8.5 embryo showing the largely mutually exclusive expression of the EC marker *Cdh5* (cyan) and *Etv2* (magenta) with the local-mesoderm markers *Six1* (green), *Meox1* (blue), *Tbx1* (red) and *Hoxc8* (yellow). Boxed regions I-IV are magnified on the right.

To examine how local mesodermal identities are remodeled during subsequent differentiation to ECs, we analyzed mesodermal, HE, and endothelial markers. Analysis of representative markers confirmed robust downregulation of local mesoderm markers such as *Six1, Meox1, Hoxc8* and *Tbx1* in *Cdh5*-positive ECs (**Figure 3D**). To further interrogate these dynamics *in situ*, we quantified expression of local mesoderm markers in discrete spatial domains across four cell states: *Kdr*-*Etv2*- mesoderm, *Kdr+Etv2−* vascular-primed mesoderm, *Kdr+Etv2*+ specified HE populations, and *Cdh5*+ committed ECs. This analysis supported the trends observed in the vascular-enriched atlas, with local mesodermal programs undergoing progressive repression during vascular specification, while a subset of markers, such as *Dlk1* and *Foxc2*, persist in committed ECs, in agreement with single cell data (**Figure S5B-C)**

### Mesodermal vascular plasticity resolves by E8.5

We next investigated the developmental plasticity of vascular-primed mesoderm using time-resolved lineage tracing. To that end, we devised an experimental framework based on the Fgd5-CreERt2 (Tamoxifen-inducible) driver mouse strain. FGD5 is involved in cytoskeletal organization downstream of VEGF signaling^45–47^, and is known to mark both blood and vascular cells^48^. In our vascular-enriched atlas, *Fgd5* is expressed in a subset of mesoderm types and at higher levels during vascular specification, while expression in blood was relatively lower than in HEs or ECs (**Figure 4A**). We therefore mated Fgd5-CreERt2 with a mouse strain harboring a ROSA26-loxP-stop-loxP-TdTomato reporter, enabling permanent genetic labeling of *Fgd5*-expressing cells and their progeny, followed by isolation of labeled cells for scRNA-seq analysis. Pregnant females were injected with 4-OHT, the active metabolite of Tamoxifen, at four time points (E6.5, E7.5, E8.5 and E9.0). Embryos induced with 4-OHT exhibited robust labeling of vasculature by E9.5, validating the specificity of the system. At this time embryos were dissected, and TdTomato-positive cells were isolated by FACS, and analyzed by scRNA-seq (**Figure 4B**). Given the transient activity of 4-OHT, the gradual increase in *Fgd5* expression in the vascular lineages and the equivalent endpoint, differences in cell type composition between the four experimental groups reflect differences in the composition of *Fgd5*-expressing cells during the temporal window of induction. Strikingly, the results indicate a “watershed” point in differentiation potential. Inductions up to E7.5 labeled not only ECs, but also a significant proportion of other embryonic mesodermal populations. In contrast, following E8.5 and E9.0 injections, labeled mesodermal derivatives were largely absent and endothelial and blood fates dominated (**Figure 4C**). These results indicate that mesoderm populations retain vascular developmental potential before E8.5, although only a subset of them will eventually differentiate into the hematoendothelial lineage. By E8.5, this broader developmental potential is largely resolved, with *Fgd5*-expressing cells contributing predominantly to vascular and hematopoietic lineages.

**Figure 4.**
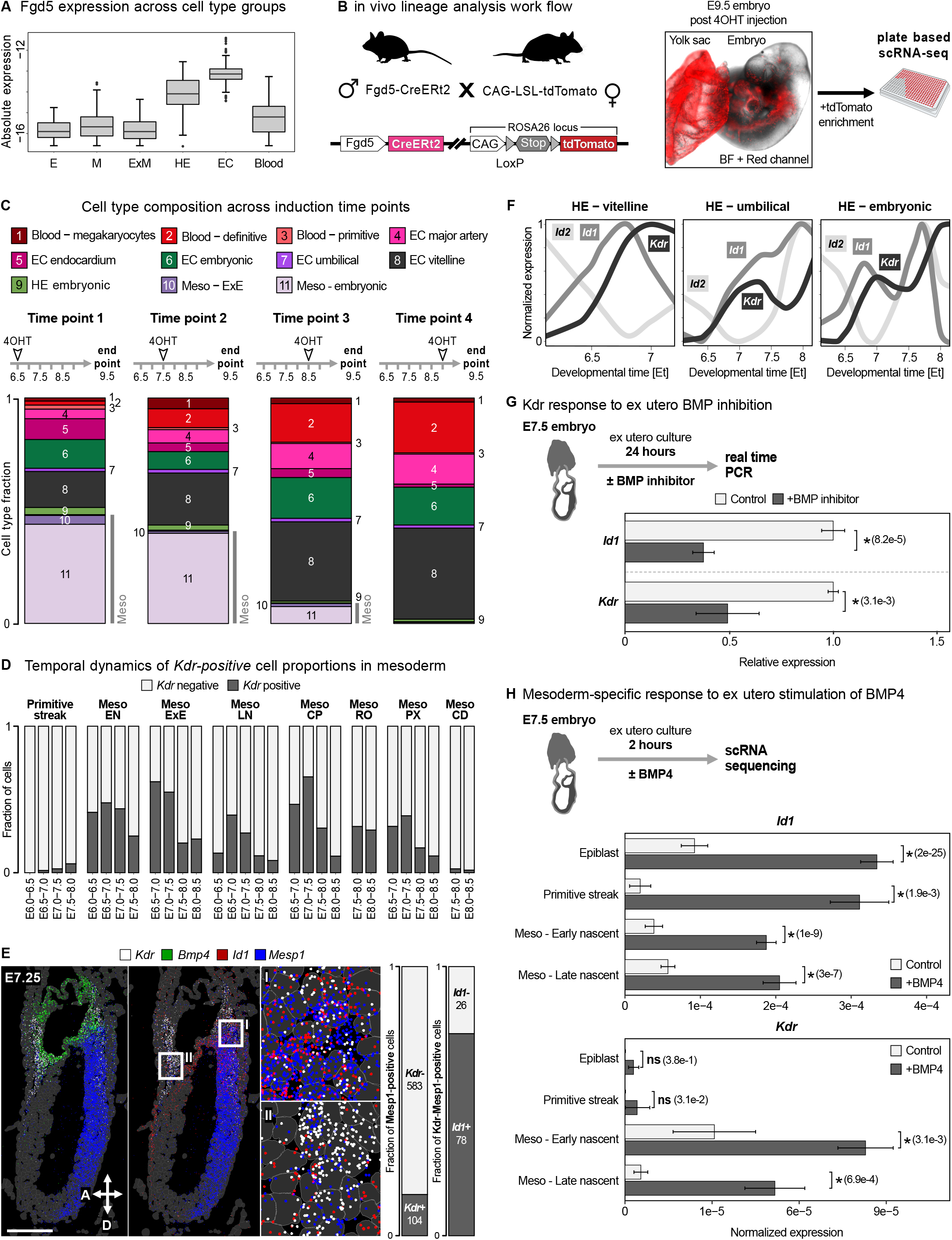
Lineage analysis reveals a window of plasticity for mesoderm specification to HE. **A,** Box plots of *Fgd5* expression across major cell type groups. **B,** Scheme of the mouse crossing strategy and experimental design for in vivo lineage tracing. Right, representative E9.5 embryo injected at E7.5. **C,** Stacked-bars of E9.5 cell type fractions from scRNA-seq of tdTomato-labeled cells across four induction time points, as indicated. **D,** Stacked-bar quantification of vascular-enriched atlas data showing the proportions of *Kdr*-negative (local mesoderm) and *Kdr*-positive (vascular-primed mesoderm) cells within each mesodermal population within 12-hour developmental intervals. **E,** MERSCOPE section of a representative E7.25 embryo showing co-localization of the BMP effector *Id1* (red) within *Kdr*-positive/*Mesp1*-positive primed mesoderm (*Kdr*, white; *Mesp1*, blue), alongside proximal *Bmp4 transcripts* (green) and *Kdr*-positive domains; boxed regions I-III are magnified on the right. Scale bar, 100 µm. Right, stacked-bar quantification of the MERSCOPE data showing the fraction of *Mesp1*-*Kdr*-positive/negative cells (left) and *Id1* in *Kdr*-positive cells (right). **F,** Normalized expression of the canonical BMP targets *Id1* and *Id2* alongside *Kdr* across developmental time (Et) in the vitelline, umbilical, and embryonic HE lineage trajectories. **G,** Real-time quantitative PCR of *Id1* and *Kdr* in *ex-utero* cultured E7.5 embryos incubated 24 hours with (n = 7) or without (n = 5) the BMP inhibitor LDN-193189. Bars show the mean; error bars, standard error of the mean; asterisk, p-value (t-test) < 0.05; ns, not significant. **H,** Single-cell RNA sequencing of ex utero cultured E7.5 embryos after 2 hours with or without BMP4, showing normalized expression of *Id1* and *Kdr* across cell types. Bars show the mean; p-values (Wilcoxon rank-sum test, Benjamini-Hochberg adjusted) are indicated in parentheses; asterisk, q-value < 0.05; ns, not significant. E, epiblast/endoderm; Meso/M, mesoderm; ExM, extraembryonic mesoderm; HE, hematoendothelial progenitors; EC, endothelial cells; EN, early nascent; LN, late nascent; CP, cardiopharyngeal; RO, rostral; PX, paraxial; CD, caudal; A, anterior; D, distal. Scale bars, 100 µm.

### BMP signaling potentiates mesoderm toward a vascular-primed state

Consistent with the lineage-tracing results, *Kdr*+*Etv2*− vascular-primed mesoderm represented a significant percentage of cells within a variety of embryonic and extraembryonic mesoderm cell types. Consistent with the flow prediction such cells are absent in the caudal mesoderm population (**Figure 4D**). Moreover, this analysis revealed a progressive decline in vascular-primed mesoderm over developmental time, mirroring the aforementioned resolution of vascular plasticity by E8.5. We next sought to identify signals associated with acquisition of the vascular-primed mesoderm state. MERSCOPE analysis of late streak (∼E7.25) embryos confirmed that *Kdr* expression is restricted to a subset of *Mesp1*-expressing early nascent mesoderm cells and further revealed that these cells are localized preferentially to the proximal region of the embryo (**Figure 4E**). A major signaling regime along the proximal-distal axis at this stage is BMP4, which is expressed early on by chorion progenitors located at the boundary between the embryo proper and extraembryonic ectoderm and later by the ExM^43,49,50^. Consistent with BMP pathway activation, *Kdr*-positive vascular-primed mesoderm cells co-express the canonical BMP target *Id1*^51^ (**Figure 4E**). Analysis of vascular trajectories further showed that *Id1/2* expression precedes or co-occurs with the induction of *Kdr* across vitelline, embryonic and umbilical HE trajectories (**Figure 4F**).

Prior genetic studies established that loss of BMP-pathway components causes early vascular defects, indicating a functional requirement for BMP in vasculogenesis^50,52^. To test whether BMP signaling is involved in potentiating mesoderm toward a vascular-primed state, E7.5 embryos were extracted and cultured ex-utero for 24 hours in the presence of the BMP inhibitor LDN-193189 followed by qRT-PCR analysis. BMP inhibition resulted in downregulation of *Kdr* expression (**Figure 4G**). We next cultured E7.5 embryos *ex utero* in the presence of BMP4 for two hours, followed by single-embryo scRNA-seq analysis of treated (n=7) compared to non-treated (n=5) time-matched embryos (**Methods**). This analysis showed that BMP4 treatment induces robust expression of *Id1* across epiblast, primitive streak, and early and late nascent mesoderm states, confirming broad competence to respond to BMP signaling. However, induction of *Kdr* was largely restricted to mesodermal populations and was not observed in epiblast or primitive streak cells (**Figure 4H** and **S6A-B**). Together, these results place BMP signaling upstream of vascular-primed mesoderm and support a model in which BMP signaling promotes vascular potentiation in a subset of mesoderm populations, at least in part through induction of *Kdr*. The incomplete repression of *Kdr* following BMP inhibition further suggests that BMP signaling is not solely responsible for establishment of the vascular-primed state. Instead, acquisition of vascular competence likely reflects integration of combinatorial signaling inputs, potentially including FGF signaling^53,54^.

### VEGF signaling is required for transition from vascular-primed mesoderm to HE states

We next investigated the signaling requirements governing the transition from vascular-primed mesoderm to specified HE states. Since *Kdr* (also known as *Vegfr2)* marks vascular-primed mesoderm, we first examined expression of its principal ligand, *Vegfa*, using the gastrulation atlas alongside MERSCOPE profiling (**Figure 5A** and **S7A**). *Vegfa* expression was detected throughout the visceral endoderm and extraembryonic visceral endoderm, which line the embryo proper and yolk-sac, respectively. Robust expression was also observed in extraembryonic mesoderm, particularly within the yolk-sac and allantois mesoderm at later stages. Beginning around E8.0, *Vegfa* expression was also detected in head mesenchyme, endothelial cells, definitive endoderm and emerging gut derivatives (**Figure 5A** and **S7B**). Thus, vascular-primed mesoderm arises in a broadly permissive VEGF signaling environment, while regions undergoing active vasculogenesis exhibit particularly high levels of *Vegfa* expression.

**Figure 5.**
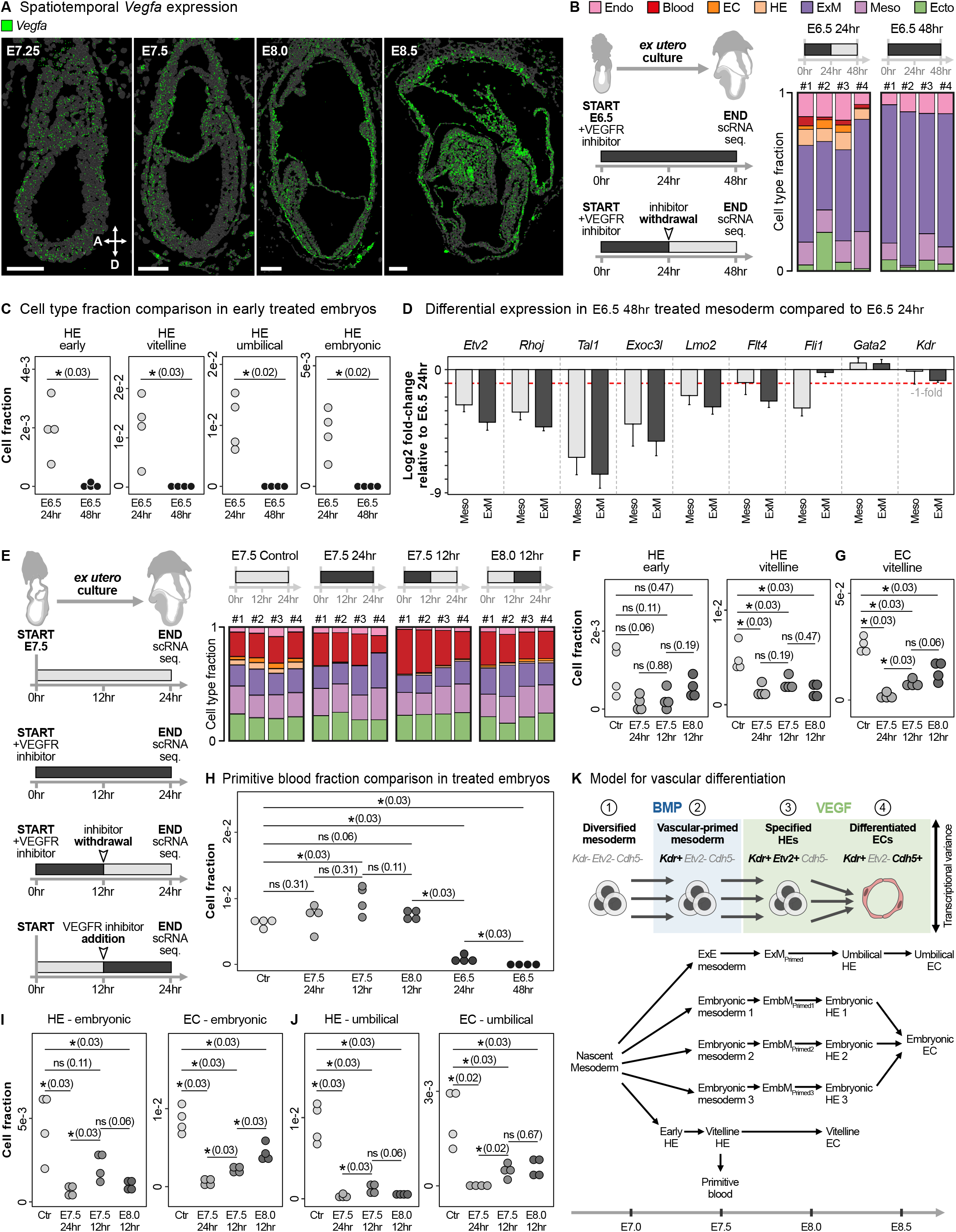
Timed perturbation of VEGF signaling reveals differential lineage-specific impact. **A,** Representative MERSCOPE sections showing *Vegfa* expression (green) from E7.25 to E8.5. Scale bars, 100 µm. A, anterior. **B,** Left, experimental scheme. Right, cell type fractions of individual embryos under 48 hours (hr) continuous VEGFR inhibition (E6.5 48hr, n = 4), or 24 hr inhibition followed by withdrawal (E6.5 24hr, n = 4). **C,** Fractions of early, vitelline, umbilical, and embryonic HE in individual embryos following 24 hr or 48 hr inhibition starting E6.5. **D,** Relative HE-module gene expression in embryonic (Meso) and extraembryonic (ExM) mesoderm, comparing 48 hr inhibition with 24 hr inhibition followed by withdrawal; the red dashed line marks a 1-fold reduction (Log2). **E,** Left, experimental scheme for (**F-J)**. Right, cell type fractions in individual control embryos, embryos under continuous 24 hr inhibition (E7.5 24hr, n = 4) and embryos pulsed with inhibitor during the first (E7.5 12hr, n = 4) or second 12 hr window (E8.0 12hr, n = 4). **F-G,** Fractions of early and vitelline HE (**F**) and vitelline EC (**G**). **H,** Primitive blood fraction in treated embryos. **I-J,** Fractions of embryonic (**I**) and umbilical (**J**) HE and EC. In (**C, F**-**J**), each dot represents one embryo. p-values (Wilcoxon rank-sum test, Benjamini-Hochberg adjusted) are indicated in parentheses; asterisk, q-value < 0.05; ns, not significant. Endo, endoderm; Blood, primitive blood; EC, endothelial cells; HE, hematoendothelial progenitors; ExM, extraembryonic mesoderm; Meso, mesoderm; Ecto, ectoderm; Ctr, control.

To test this hypothesis, we cultured gastrulation-stage embryos ex utero in the presence of Axitinib, a selective VEGFR inhibitor^55,56^. Embryos extracted at E6.5 were exposed to VEGFR inhibition under one of two regimes: continuous inhibition for 48 hours, or transient inhibition for the first 24 hours followed by withdrawal and culture for another 24 hours. All embryos were subsequently profiled individually by scRNA-seq (**Methods**, **Figure 5B**). Continuous VEGFR inhibition completely abolished hematoendothelial derivatives, including HE, endothelial and blood populations. This is consistent with hematopoietic and endothelial defects previously reported in *Kdr* knockout mice^57^. Strikingly, when the inhibitor was withdrawn after 24 hours, early, vitelline, embryonic and umbilical HE populations, together with their endothelial derivatives, re-emerged (**Figure 5C** and **S7C**). We next asked whether VEGFR inhibition affects the vascular-primed state itself or the subsequent transition toward HE specification. To address this, we compared gene expression in pooled embryonic and extraembryonic mesodermal populations from embryos subjected to continuous VEGFR inhibition with embryos in which the inhibitor was withdrawn after 24 hours. Expression of HE-module genes, including *Etv2*, *Rhoj* and *Tal1*, was strongly reduced following continuous inhibition, but recovered after withdrawal. In contrast, *Kdr* expression was only modestly affected (**Figure 5D**). These results support the notion that VEGF signaling is not required for mesodermal acquisition of a vascular-primed state but is required for the subsequent transition into specified HE progenitors.

### VEGF signaling modulates hematoendothelial outcomes from early and vitelline HEs

Although the experiments above establishes a requirement for VEGF signaling during the transition from vascular-primed mesoderm to specified HE progenitors, they do not resolve whether all HE trajectories exhibit similar temporal requirements for VEGF signaling. Furthermore, prolonged ex utero culture altered the relative composition of embryonic cell states compared with stage-matched non-cultured embryos. We therefore performed a shorter perturbation analysis of 24 hours from E7.5, a condition in which overall cell-state composition more closely resembles WT embryos. To examine the temporal requirements for VEGF signaling across individual HE trajectories, embryos were exposed to VEGFR inhibition under one of four regimes: untreated control, continuous inhibition for 24 hours, inhibition during the first 12 hours of culture followed by withdrawal, or inhibition during the final 12 hours of culture following an initial 12 hours without inhibition. (**Figure 5E**).

We first analyzed the impact of temporal VEGFR inhibition on early and vitelline HEs, which represent the earliest HE populations (**Figure 5F**). Early HEs are specified predominantly between E7.0 and E7.5 and persist only transiently thereafter, whereas vitelline HEs emerge from E7.5 onward (**Figure 1I**). Because the perturbation experiments were initiated at E7.5 and analyzed 24 hours later, only limited numbers of early HEs were expected. Consistent with this expectation, early HEs were detected at low abundance across all perturbation conditions and exhibited little variation between treatments. Vitelline HEs, by contrast, were strongly affected by the timing of VEGFR inhibition. Continuous inhibition throughout the culture period resulted in a marked reduction of vitelline HEs relative to untreated controls, consistent with their specification during this developmental interval. Restricting inhibition to the first 12 hours of culture followed by inhibitor withdrawal, partially restored vitelline HE abundance. Surprisingly, delaying inhibition until the second 12-hour interval reduced vitelline HE numbers to levels comparable with continuous inhibition (**Figure 5F**).

Analysis of vitelline ECs, which arise from vitelline HEs, provided additional insight into these dynamics. Continuous VEGFR inhibition resulted in a marked reduction of vitelline ECs, whereas inhibition limited to the first 12 hours was associated with partial rescue (**Figure 5G**). This pattern broadly mirrors the corresponding changes observed in their vitelline HE progenitors. In contrast, delaying inhibition until the second 12-hour interval resulted in a further increase in the numbers of vitelline ECs despite reduced numbers of vitelline HEs. One possible explanation is that vitelline HEs generated during the first 12 hours had already initiated endothelial differentiation prior to inhibitor treatment. Consequently, fewer cells remained in the HE state at the endpoint, while larger numbers accumulated as differentiated ECs.

Given the proposed contribution of early and vitelline HEs to primitive hematopoiesis, we next examined the effects of VEGFR inhibition on primitive blood. While continuous VEGFR inhibition from E6.5 throughout the 48-hour culture period completely abolished primitive blood, transient inhibition during the first 24 hours of culture followed by inhibitor withdrawal still permitted primitive blood formation, albeit at very low levels. In contrast, all perturbations initiated at E7.5 onward resulted in primitive blood levels comparable to control embryos (**Figure 5H**). Since early HEs are specified predominantly before E7.5, this pattern is consistent with a contribution of early HEs to primitive hematopoiesis. Interestingly, primitive blood abundance nevertheless varied among the E7.5 perturbation conditions. Primitive blood was modestly increased following continuous inhibition and further increased when inhibition was restricted to the first 12 hours of culture, whereas delaying inhibition until the second 12-hour interval reduced primitive blood abundance to levels comparable with untreated controls. This pattern broadly mirrors the corresponding changes observed in vitelline HEs and is therefore consistent with an additional contribution of vitelline HEs to primitive hematopoiesis. Notably, the reduction in primitive blood under delayed inhibition coincided with increased abundance of vitelline ECs (**Figure 5G**), suggesting that VEGF signaling may promote EC differentiation at the expense of primitive blood within the vitelline trajectory.

### Temporal VEGF effects follow embryonic and umbilical HE specification windows

Embryonic and umbilical trajectories exhibited responses that reflect the developmental intervals during which these populations are specified (**Figure 1I**). Embryonic HEs are generated predominantly between E7.5-8.5 and exhibited a pattern similar to that observed in the vitelline trajectory (**Figure 5I**). Continuous VEGFR inhibition resulted in robust repression of embryonic HEs and their corresponding endothelial derivatives, whereas inhibition restricted to the first 12 hours followed by inhibitor withdrawal increased both populations. In contrast, delaying inhibition until the second 12-hour interval reduced embryonic HE abundance while producing the largest accumulation of embryonic ECs, consistent with endothelial differentiation of embryonic HEs prior to inhibitor treatment. Umbilical HEs, which emerge predominantly after E8.0, remained strongly repressed under all inhibition regimes, consistent with specification throughout the entire perturbation window. Their corresponding endothelial derivatives exhibited a similar response (**Figure 5J**). Finally, differential expression analysis showed that HEs recovering in each treatment group closely resembled control HEs transcriptionally, indicating that perturbation alters the timing of their emergence without compromising their identity, and further supporting a model in which HE populations are continuously recruited from distinct vascular-primed mesoderm states (**Figure S7D**).

## DISCUSSION

The close developmental relationship between blood and endothelial cells was first recognized through observations of yolk-sac blood islands, where primitive blood cells arise in association with the surrounding vascular endothelium. This observation led to the proposal that both lineages may arise from common progenitors. The concept was formalized through the haemangioblast model and later refined by the discovery of hemogenic endothelium as an intermediate stage in definitive blood development^10,11,17^. In parallel, genetic studies established VEGF signaling as an essential regulator of vasculogenesis^58^, with Vegfr2/*Flk1*/*Kdr* marking early vascular progenitors, and loss of *Kdr* resulting in profound defects in blood-island formation and vascular development^57,59^. Clonal analysis subsequently demonstrated that individual yolk-sac blood islands arise from multiple progenitors rather than a single founder cell^5^, while recent lineage tracing identified evidence that yolk-sac hematopoiesis cannot be explained solely by a classical hemangioblast model^4^. Despite these insights, the developmental process itself has remained difficult to reconstruct. In particular, how vascular progenitors emerge from gastrulating mesoderm, when endothelial specification occurs, how distinct vascular populations are related to one another, and where VEGF signaling acts within this sequence have remained unresolved.

Here, we reconstruct vasculogenesis as a continuous developmental process spanning embryonic and extraembryonic tissues (**Figure 5K**). This framework reveals a sequential developmental program that begins shortly after mesodermal cells exit the primitive streak. A subset of these cells first enters a transient *Kdr*+*Etv2*− vascular-primed state before activating the vascular specification program. BMP4 signaling contributes to the establishment of this state, as BMP4 stimulation promotes *Kdr* expression, whereas inhibition of BMP signaling attenuates it. Additional posterior signaling pathways, including FGF, WNT and BMP2, may also contribute to this process^60–62^. Importantly, the vascular-primed state precedes blood and vascular commitment but does not precede mesoderm diversification, precluding the possibility of a common progenitor for all vascular cells. Thus, regional mesodermal identities are first established, and only subsequently do cells activate *Etv2* and associated vascular regulators, including *Tal1*, *Lmo2* and *Hhex*, followed by endothelial markers such as *Cdh5*, *Pecam1* and *Cd93*. *Etv2* expression is strongly associated with the hemato-endothelial cell state, often considered equivalent to the classically proposed hemangioblast - a committed, bipotent progenitor. However, this state differs in two critical ways: first, *Etv2*-expressing lineages possess a broader developmental capacity than endothelium and blood^63^. Second, HEs are defined as a stage within a cell-state trajectory rather than a cellular lineage and, as such, do not dictate a rigid cell-fate prediction. Together, these observations support a developmental model in which mesoderm diversification precedes vascular specification. Lineage-tracing experiments further suggest that this vascular-primed state persists until approximately E8.0-8.5, after which transitional populations largely resolve into committed endothelial and local mesodermal derivatives. Alongside this, a phase of post-gastrulation vasculogenesis is also known to contribute significantly to nascent vasculature^35–42^. The reconstructed trajectories further suggest that hematoendothelial potential is not uniformly distributed across vascular populations. Instead, dual hematoendothelial potential appears largely restricted to the earliest trajectories, namely Early and vitelline HEs, whereas later embryonic and umbilical endothelial trajectories contribute predominantly to endothelial lineages.

Our perturbation experiments identify VEGF signaling as an essential requirement for vascular specification from vascular-primed mesoderm. VEGFR inhibition arrests vasculogenesis, preventing the emergence of endothelial trajectories while preserving the underlying vascular-primed mesodermal population. Notably, VEGFR inhibition prevents the specification of both primitive blood and endothelial lineages, indicating that VEGF signaling acts before their divergence. Upon release from inhibition, endothelial differentiation resumes from this pre-existing primed state. These findings place VEGF signaling downstream of mesoderm diversification and vascular priming, but upstream of vascular and hematopoietic specification, independently validating the developmental sequence inferred from the reconstructed trajectories. Temporal perturbation experiments further reveal a later role for VEGF signaling following hematoendothelial specification. Within the vitelline hematoendothelial trajectory, VEGF signaling appears to balance blood and endothelial cell production in favor of the latter, indicating that VEGF contributes both to the specification of vascular progenitors and to subsequent endothelial differentiation. Although our experiments do not distinguish between individual VEGF ligands, VEGFA is likely to be the principal mediator of these effects in the early embryo.

Finally, our findings reveal that, in contrast to the progressive diversification that characterizes most developmental trajectories, vascular populations arising from distinct embryonic and extraembryonic mesodermal origins converge toward a common transcriptional program. Similar convergence has been described during differentiation of neuromesodermal progenitors toward neural and mesodermal derivatives^7,30,64,65^ and during the merging of visceral and definitive endoderm into the gut endoderm^66^. Although much of the transcriptional signature associated with their mesodermal origins is progressively lost during differentiation, remnants of this developmental history persist and may help explain the developmental origins of organotypic endothelial identities described later in development^8,18–20^.

## METHODS

### Embryo recovery and animal procedures

All animal procedures were approved by the Institutional Animal Care and Use Committee and were performed in strict adherence to Weizmann Institute guidelines. Mice were monitored for health and activity and were given ad libitum access to water and standard mouse chow with 12-hr light/dark cycles. Embryos were collected from timed pregnant C57BL/6JRccHsd, Hsd:ICR(CD-1) or B6;129S6-Gt(ROSA)26Sortm9(CAG-tdTomato)Hze/J (JAX strain 007905) females (obtained from Envigo or JAX and mated in house with males of the same strain), between E6.5-9.5.

### Immunostaining

In this study immunostaining was performed following standard protocols. The following primary antibodies were used in a 1:100 ratio: Goat anti-mouse VEGFR2/KDR/Flk-1 (AF644, R&D Systems); Goat anti-mouse LYVE-1 (AF2125, R&D Systems); Goat anti-mouse Pref-1/DLK1/FA1 (AF8272, R&D Systems). Secondary antibodies were purchased from Jackson ImmunoResearch, and used in 1:200 ratio.

### Flow Cytometry

For isolation of single cells for scRNA-seq, embryos were dissociated with 0.25% Trypsin-A, 0.02% EDTA (Biological Industries) solution for 5 minutes at 37℃, and re-suspended in DMEM without phenol red (GIBCO) supplemented with 10% FBS (Biological Industries). Samples were run on a FACS Aria-III flow cytometer (BD Biosciences, using BD FACSDiva Software v9.0) using the ‘index sort’ option to retain the spectral properties of each individual sorted cell. For spatial-informed samples (Figure 1F) obtained, we further dissected regions of interest (described in text). Subsequently, we employed index-sorting and enriched for KDR-positive cells using Rat anti-mouse KDR antibody (136403, BioLegend) in a 1:100 ratio.

### Lineage analysis using Fgd5-cre/ERT2 mice

Timed pregnancies were generated by mating B6;129S6-Gt(ROSA)26Sortm9(CAG-tdTomato)Hze/J females with C57BL/6N-Fgd5tm3(cre/ERT2)Djr/J males (JAX strain 027789). Pregnant females were administered 4-hydroxytamoxifen (4-OHT; 50mg/kg) at four time points as indicated in Figure 4 through intraperitoneal injection. Embryos were collected at E9.5 and processed for single-cell RNA sequencing as described below (MARS-seq).

### Signaling perturbations of ex utero cultured post-implantation embryos

Embryos were dissected either at E6.5 or E7.5, their Reichert’s membrane removed and up to 3 embryos were placed into separate wells of 8-well glass bottom/ibiTreat μ-plates (iBidi; 80827/80826) filled with 250μl of EUCM (consisted of 25% DMEM (GIBCO 11880; includes 1mg/mL D-glucose and pyruvate, without phenol red and without L-glutamine) supplemented with 1x Glutamax (GIBCO, 35050061), 100 units/ml penicillin/100 μg/ml streptomycin (Biological industries; 030311B) and 11 mM HEPES (GIBCO, 15630056), plus 50% Rat Serum (RS) (Rat whole embryo culture serum, BIOIVT) using the same protocol as in Aguilera-Castrejon et al., 2021. For two hours BMP4 induction experiments (Figure 4F and S6), EUCM did not contained RS in order to control for other potential signaling molecules in media. In all experiments media was pre-heated and gas calibrated for half an hour in an incubator with 5% CO2 at 37°C. Embryos were cultured statically for 2, 24 or 48 hours as indicated in the text in an incubator with 5% CO2 at 37°C. EUCM was supplemented with either 200ng/μl Axitinib (R&D systems, 4350), 1000ng/mL BMP4 (R&D, 5020-BP), or 5µM LDN-193189 (Sigma-Aldrich, SML0559). After incubation embryos were collected and processed for qRT-PCR (Figure 4G) or single-cell RNA sequencing (Figure 4H and S6, Figure 5B-J and S7D) as described below.

### MERSCOPE spatial transcriptomics

Spatial gene expression profiling was performed on 7 embryos at E7.5-8.5 using the MERSCOPE V2 platform (Vizgen). A custom panel of 500 genes was designed based on differential expression analysis across our wildtype single-cell atlas^43^, selecting genes with the most robust transcriptional specificity across cell populations. Tissue sections of 4µm were standardly paraffinized from pre-fixed embryos (4% PFA) in 2% ultra-low melting agarose and processed according to the manufacturer’s protocol. Imaging was performed using the MERSCOPE Ultra Spatial Imaging Platform. Images were further segmented into individual embryos using the MERSCOPE Vizualizer Software (Figure 2D-F, S3, Figure 4E, and S5B). Cell segmentation was performed using Cellpose 2.0 cell boundary kit^67^.

### Quantification of MERSCOPE results

Cell segmentation and manual per-embryo assignment (above) produced, for each section, a cell-by-gene transcript count matrix and a matched per-cell metadata table containing the cell segmentation identifiers, spatial coordinates and embryo annotations. For each analysis, this metadata was used to retain the cells belonging to the embryos or regions of interest and the count matrix was subset to these cells accordingly. Per-cell transcript counts were used directly for thresholding, or normalized to within-cell fractions by dividing each cell by its total transcript count where relative expression was required. To classify cells by marker status (Figure 2F and Figure 4E), cells were considered positive for a gene when transcript count exceeded a gene-specific detection threshold, defined as the 50th percentile of that gene’s counts across the cell population, thereby adapting to each gene’s dynamic range and detection level. Cells were then assigned to co-expression categories (e.g. double-negative, single-positive or double-positive), and the fraction of cells in each category was computed within the relevant marker-positive population. To assess how regional mesodermal identity relates to endothelial commitment, marker-positive cells within a defined region were further stratified by their endothelial differentiation status (Figure S5B). Using the E8.5 Cdh5 MERSCOPE dataset, cells were partitioned by region (head and trunk; tail and yolk sac) from the manual annotations, and region-specific mesodermal identity markers were examined (*Six1*, *Meox1*, *Tbx1, Msx1* and *Foxc2* in the head and trunk; *Hoxc8* and *Dlk1* in the tail and yolk sac). For each gene, positivity was called using a threshold set at the 50th percentile of its counts across the cell population, applied to both the mesodermal markers and the endothelial genes (*Kdr*, *Etv2* and *Cdh5*). Marker-positive cells were assigned to an ordered series of progressively committed endothelial states: marker-positive but negative for *Kdr*, *Etv2* and *Cdh5*; additionally *Kdr*-positive; additionally *Etv2*-positive; and *Cdh5*-positive. The mean ± s.e.m. of normalized marker expression was computed for each state. Expression in each state was compared to the marker-positive, endothelial-negative baseline using two-sided Wilcoxon rank-sum tests, with Benjamini-Hochberg correction across states. For each gene, expression levels were compared between cells from different populations.

### Single-cell RNA sequencing

#### 10x Genomics

Cultured embryos were prepared for sequencing at different developmental stages. Embryos were selected for each developmental stage, pooled and dissociated using Trypsin-EDTA solution A 0.25% (Biological Industries; 030501B) for 5 minutes at 37℃. Trypsin was neutralized with media including 10% FBS and cells were washed and re-suspended in 1x PBS (calcium and magnesium free) with 400μg/ml BSA. Cell suspension was next filtered with a 100μm cell strainer to avoid cell clumps. Single cell viability was determined by trypan blue staining, before being diluted to a final concentration of 700-1600cells/μl. Single-cell RNA-seq libraries were generated for individual embryos using GEM-X Universal 3’ Gene Expression v4 4-plex (5000 cell recovery target) and sequenced on the AVITI System by Element Biosciences according to the manufacturer’s instructions.

#### MARS-seq

Single-cell cDNA libraries were prepared as previously reported^68^ following the MARS-seq protocol^69^.

### Single-cell RNA sequencing data processing

#### 10x Genomics data from ex utero cultured embryos

Raw files were transformed into count matrices using Cell Ranger 9.0.1 with default parameters and with the pre-built Cell Ranger reference package refdata-gex-mm10-2020-A (mm10 genome, GENCODE vM23/Ensemble 98). Cells with less than 1000 counts, more than 20000 counts or a high number of counts from mitochondrial genes relative to the number of counts from ribosomal genes were removed.

#### ^43^MARS-seq

Libraries were sequenced using NovaSeq 6000. Reads were processed following the MARS-seq2.0 protocol^69^ with the same specifications as previously reported^43^ using the Cell Ranger 9.0.1 with default parameters and with the pre-built Cell Ranger reference package refdata-gex-mm10-2020-A (mm10 genome, GENCODE vM23/Ensemble 98).

### Vascular-enriched atlas construction

To build the vascular atlas, we integrated cells from all experiments performed on E7.5-9.0 embryos, including spatially informed KDR-enriched and lineage analysis datasets, together with wildtype cells previously annotated as mesoderm and extraembryonic mesoderm subtypes^43^ from embryos dissected between E6.5-8.5. The combined dataset was processed using the Metacell-2 pipeline^70^, retaining genes that met a minimum variance-to-mean ratio and coverage threshold. MetaCell (MC) clusters enriched for cell-cycle- or stress-related gene signatures were manually identified and removed. The final MC model (minimum MC size = 20) is comprised of 1,656 MCs derived from 46,265 cells. The resulting MC object was projected onto the previously published atlas^43^, and vascular subtypes were manually annotated based on gene signature (Figure S1A and Supplementary Information Table 1) and spatiotemporal information (Figure 1F). A separate MC object for ex utero-cultured embryos was constructed using the same feature genes and parameters.

### Vascular atlas annotation refinement

Initial HE and EC subtype annotations were assigned based on the vascular atlas MC structure and spatiotemporal information. To refine annotation boundaries and correct metacells whose gene expression profiles were more consistent with a neighboring cell type than their initially assigned one, we applied an iterative correlation-based reassignment procedure separately for HE and EC subtypes. For each subtype group, a mean log-normalized expression profile was computed across the marker genes of that type using all metacells initially assigned to it. Each MC *m* was then correlated (Pearson) against all cell type mean profiles over the marker gene set 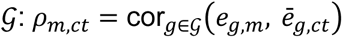 where *e_g_*_,*m*_ is the log-normalized expression of gene *g* in MC *m*, and *e̅_g_*_,*ct*_ is the mean expression across all metacells of cell type *ct*. A MC was reassigned from its original cell type *ct*_0_ to the best-correlated cell type *ct*^∗^ = arg max*_ct_ ρ_m_*_,*ct*_ if its correlation to its own cell type fell below a threshold 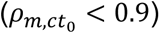 and *ct*^∗^ ≠ *ct*_0_. This procedure was applied first to HE subtypes, then to EC subtypes using the updated annotations from the previous step. The final refined annotations were used for all downstream analyses.

### Network flow model construction

We previously introduced a network flow model to reconstruct cellular differentiation trajectories during mouse gastrulation from single-embryo, single-cell transcriptomic data^30,43^. As this model requires uniform cell sampling across the atlas, trajectory inference on the extended vascular atlas was restricted to the sampled wildtype cells, excluding the enriched vascular cells that violate this assumption. The network flow model was otherwise computed as previously described^30,43^.

### Gene-module scores

#### Cell type/functional scores

Given a list of genes (referred to as "Cell-state markers") *Gm*, the single-cell score corresponding to a particular cell is calculated as the sum of counts for all genes in a given cell-state markers *Gm*, divided by the total sum of expression counts for all genes in the cell. Artery and Cdkn scores were calculated as described, arterial marker genes are *Efnb2, Dll4, Notch1, Notch4, Hey1, Hey2, Nrp1, Gja5*, Cdkn marker genes are *Cdkn1a*, *Cdkn1b*, *Cdkn1c*.

#### Cell-cycle/proliferation scores

Calculated with the same parameters and gene markers as previously described^43^.

### Atlas projection and cell type annotation of cells from ex utero cultured embryos

Following the Metacell-2 pipeline described above, each cell from *ex utero* cultured embryos was assigned a coarse cell type annotation (i.e., without subdivision into vascular subtypes). Cells annotated as haematoendothelial progenitors (HE), endothelial cells (EC), or erythroid cells were then projected onto the vascular atlas using a correlation-based matching procedure.

Let *u_g_*_,*c*_ denote the UMI count of gene *g* in query cell *c*, and *e_g_*_,*m*_ the aggregated UMI count of gene *g* in reference MC *m*. To account for differences in sequencing depth, all cells were downsampled to a common library size *d*, defined as:

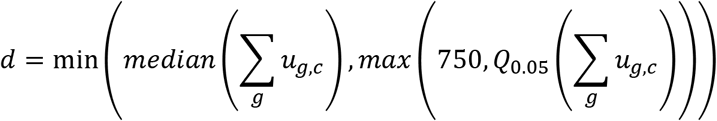

with cells having fewer than *d*total UMIs retained without downsampling. Normalized log-expression profiles were then computed as:

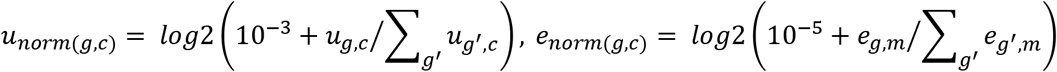

For each query cell *c*, Pearson correlations were computed over the feature gene set G against all reference metacells:

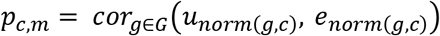

Each cell *c* was assigned the cell type label of the reference MC arg max *m p_c_*_,*m*_. Cells outside the HE/EC/erythroid lineages retained their coarse annotation from the initial Metacell-2 projection.

### Cell type frequency comparison between ex-utero-cultured embryos

For each embryo, cell type frequencies were computed as the fraction of cells assigned to each annotated cell type out of the total number of cells from that embryo. Cell types with a mean frequency below 10^−4^ across all embryos were excluded. To compare cell type composition between ex utero cultured embryos and controls, pairwise Wilcoxon rank-sum tests were performed for each cell type independently, comparing each treatment group against the control group across individual embryos (*n* = 4 per group, Figure 5). Multiple testing correction was applied across all cell types within each pairwise comparison using the Benjamini-Hochberg procedure (via q-value with *π*_0_ = 1). Results are reported as q-values, with *q* < 0.05 considered statistically significant.

### Differential expression analysis for *ex utero*-cultured embryos

Individual BMP4 treated embryos (*n* = 7) were assigned a best-matching WT rank as previously reported^43^. Cells were projected as described above, and analyses were restricted to the cell types of interest including epiblast, primitive streak, and early and late nascent mesoderm. For each comparison, the raw UMI count matrix was downsampled to a uniform depth as described above to control for differences in cell coverage. Downsampled counts were converted to within-cell expression fractions by normalizing each cell to its total UMI content. Expression of gene markers was quantified per cell as the normalized UMI fraction and summarized as the mean ± s.e.m. across cells for each cell type and treatment. Comparisons between BMP4 and stage-matched WT groups were performed using two-sided Wilcoxon rank-sum tests, computed independently for each gene and cell type. P values were corrected for multiple testing across cell types within each gene using the Benjamini-Hochberg procedure, and an adjusted P < 0.05 was considered significant. To treat embryos as biological replicates and account for embryo-to-embryo variability, the analysis was repeated at the embryo level. For each gene, per-cell normalized expression was averaged within each embryo across the nascent mesoderm compartment (early and late nascent mesoderm), retaining only embryos contributing at least 10 cells, and the resulting per-embryo means were compared between BMP4 and WT-matched embryos by two-sided Wilcoxon rank-sum test with Benjamini-Hochberg correction. For Figure 5d differential gene expression (DGE) analysis was performed using a pseudobulk approach with DESeq2^71^. Cells were first grouped into broad cell type categories, mesoderm (M) and extraembryonic mesoderm (ExM), and DGE was run both at the level of these broader groups. For each cell type group and pairwise treatment comparison, UMI counts were aggregated per embryo to generate pseudobulk profiles, retaining only embryos contributing at least 20 cells to the relevant group. Comparisons were further restricted to treatment pairs for which at least two embryos per condition were available. The DESeq2 model was fit with treatment as the sole covariate, and size factors were estimated internally by DESeq2 using its default median-of-ratios normalization. Results were visualized as log_2_ fold change, with a predefined HE gene module including *Etv2*, *Kdr*, *Gata2*, *Fli1*, *Tal1*, *Rhoj*, *Flt4*, *Lmo2*, *Exoc3l*.

## ACKNOWLEDGEMENTS

We would like to thank the Stelzer lab members for comments and advice in the preparation of the manuscript. Yosef Addadi, for imaging consultation. Y.S. is the incumbent of the Louis and Ida Rich Career Development Chair and is supported by European Research Council (ERC_CoG EmbryoCellEnsemble), ISF (2824/24), the Minerva Foundation, Helen and Martin Kimmel Stem Cell Institute. This research was generously supported by the EKARD Research School of Biological Science, John L. and Vera Schwartz, the Abisch-Frenkel Foundation for the Promotion of Life Sciences, the Dr. Dvora and Haim Teitelbaum Endowment Fund, and Magnus Konow in Honor of his mother Olga Konow Rappaport. K.Y. is the incumbent of the Enid Barden and Aaron J. Jade Professorial Chair in Memory of Canter John Y. Jade and is supported by the Horizon Europe ERC-2024-SyG (BEMOSAIC 101167365).

## AUTHOR CONTRIBUTIONS

R.B.Y., H.R., and Y.S. conceived and designed the experiments and performed data analysis, interpretations and wrote the manuscript with input from all authors. S.G. under the supervision of H.A. and S.C. assisted with data collection. N.R. prepared scRNA-seq libraries. A.H.O. assisted with animal handling and related procedures. Y.M., T.S., L.A., and D.H. assisted with MERSCOPE data collection. K.Y. contributed to the interpretation of the results and manuscript preparation. A.T. contributed and advised on computational analysis.

## ADDITIONAL INFORMATION

Supplementary Information is available for this paper. Correspondence and requests for materials should be addressed to Y.S. at.

## DATA AVAILABILITY

All sequencing data that underpin the conclusions of this study have been meticulously archived and are publicly accessible through the National Center for Biotechnology Information Gene Expression Omnibus (GEO). These data are cataloged under the GEO Series accession number GSE348021 and can be accessed directly via the following link: https://www.ncbi.nlm.nih.gov/geo/query/acc.cgi?acc=GSE348021. This ensures comprehensive availability and transparency of the data supporting our research findings.

## CODE AVAILABILITY

All custom scripts used in this study have been made openly accessible and can be found at https://github.com/hernanRubinstein/vascular-specification. Additionally, these scripts have been deposited for permanent archiving and citation at https://doi.org/10.5281/zenodo.21202838, ensuring transparency and reproducibility of the computational methods employed in this work.

