## Supplementary Figures for "Vascular specification from diversified mesoderm culminates in endothelial convergence"

### Figure S1

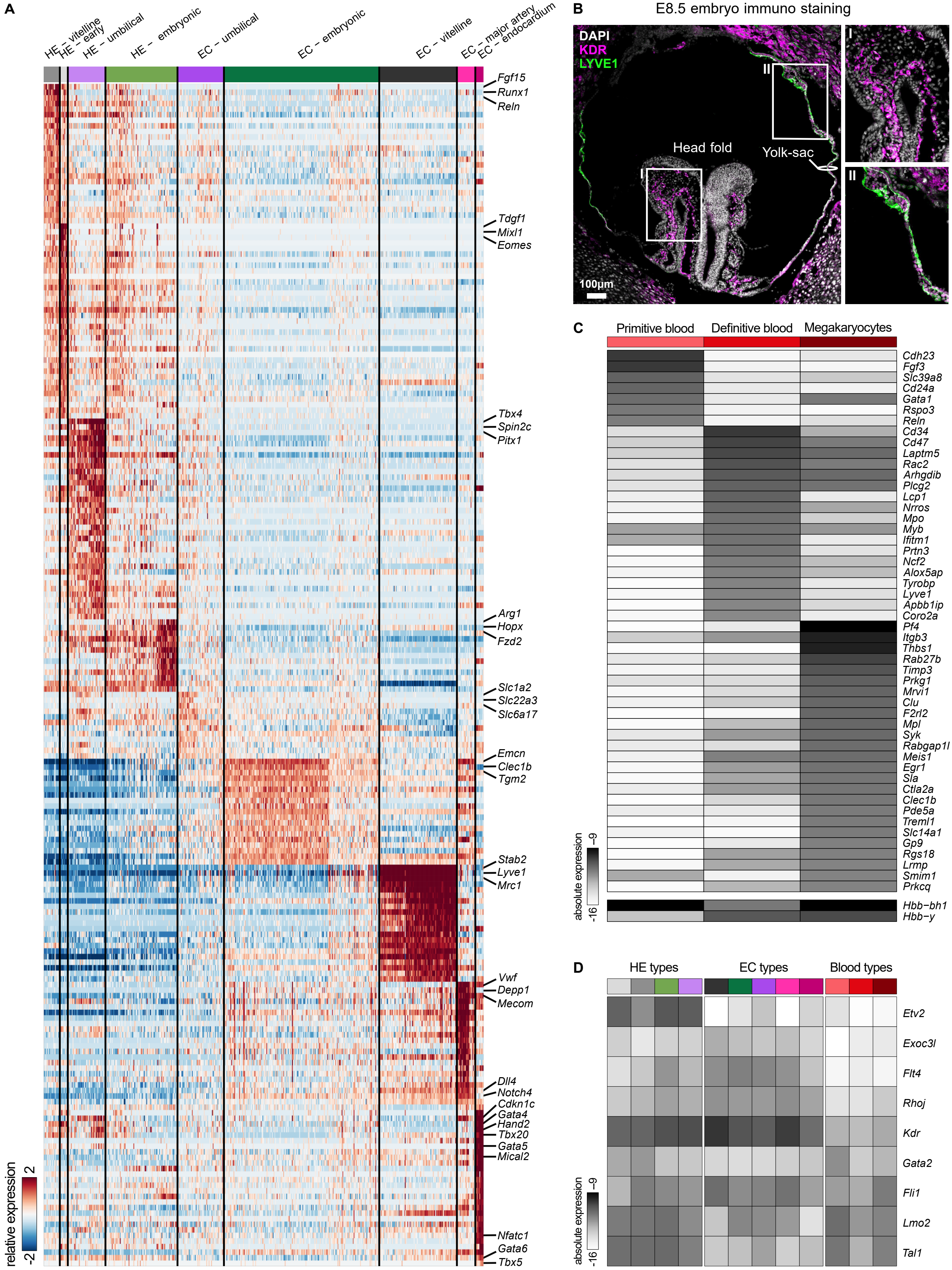
A
HE – vitelline
HE – early
HE – umbilical
HE – embryonic
EC – umbilical
EC – embryonic
EC – vitelline
EC – major artery
EC – endocardium
Fgf15
Runx1
Reln
Tdgf1
Mixl1
Eomes
Tbx4
Spin2c
Pitx1
Arg1
Hopx
Fzd2
Slc1a2
Slc22a3
Slc6a17
Emcn
Clec1b
Tgm2
Stab2
Lyve1
Mrc1
Vwf
Depp1
Mecom
Dll4
Notch4
Cdkn1c
Gata4
Hand2
Tbx20
Gata5
Mical2
Nfatc1
Gata6
Tbx5
relative expression
2
-2
B
E8.5 embryo immuno staining
DAPI
KDR
LYVE1
Head fold
Yolk-sac
I
II
100µm
C
Primitive blood
Definitive blood
Megakaryocytes
Cdh23
Fgf3
Slc39a8
Cd24a
Gata1
Rspo3
Reln
Cd34
Cd47
Laptm5
Rac2
Arhgdib
Plcg2
Lcp1
Nrros
Mpo
Myb
Ifitm1
Prtn3
Ncf2
Alox5ap
Tyrobp
Lyve1
Apbb1ip
Coro2a
Pf4
Itgb3
Thbs1
Rab27b
Timp3
Prkg1
Mrvi1
Clu
F2rl2
Mpl
Syk
Rabgap1l
Meis1
Egr1
Sla
Ctla2a
Clec1b
Pde5a
Treml1
Slc14a1
Gp9
Rgs18
Lrmp
Smim1
Prkcq
Hbb-bh1
Hbb-y
absolute expression
-9
-16
D
HE types
EC types
Blood types
Etv2
Exoc3l
Flt4
Rhoj
Kdr
Gata2
Fli1
Lmo2
Tal1
absolute expression
-9
-16

**Figure S1 | A,** Heatmap of differentially expressed genes across the HE and EC subtypes; color, relative expression. Selected markers are labeled. **B,** Immunostaining of an E8.5 embryo section for KDR (magenta) and LYVE1 (green) with DAPI (white); boxed regions I and II are magnified on the right. Scale bar, 100 μm. **C,** Heatmap of marker genes absolute expression across the blood subtypes. **D,** Heatmap of HE-module gene absolute expression across the HE, EC and blood subtypes. HE, hematoendothelial progenitors; EC, endothelial cell.

### Figure S2

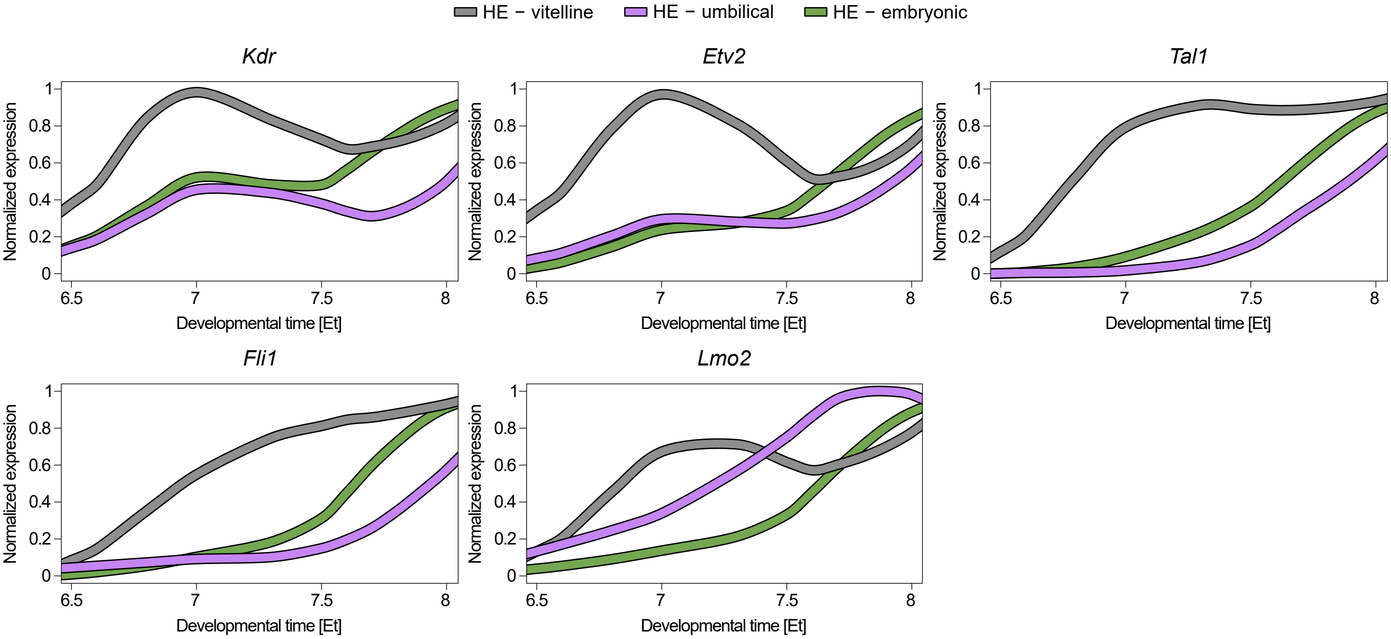

HE − vitelline
HE − umbilical
HE − embryonic
Kdr
Etv2
Tal1
Fli1
Lmo2
Normalized expression
Developmental time [Et]
0
0.2
0.4
0.6
0.8
1
6.5
7
7.5
8

**Figure S2 |** Normalized expression line plots of individual HE-module genes across developmental time (Et), in the vitelline, umbilical and embryonic HE lineages.

### Figure S3

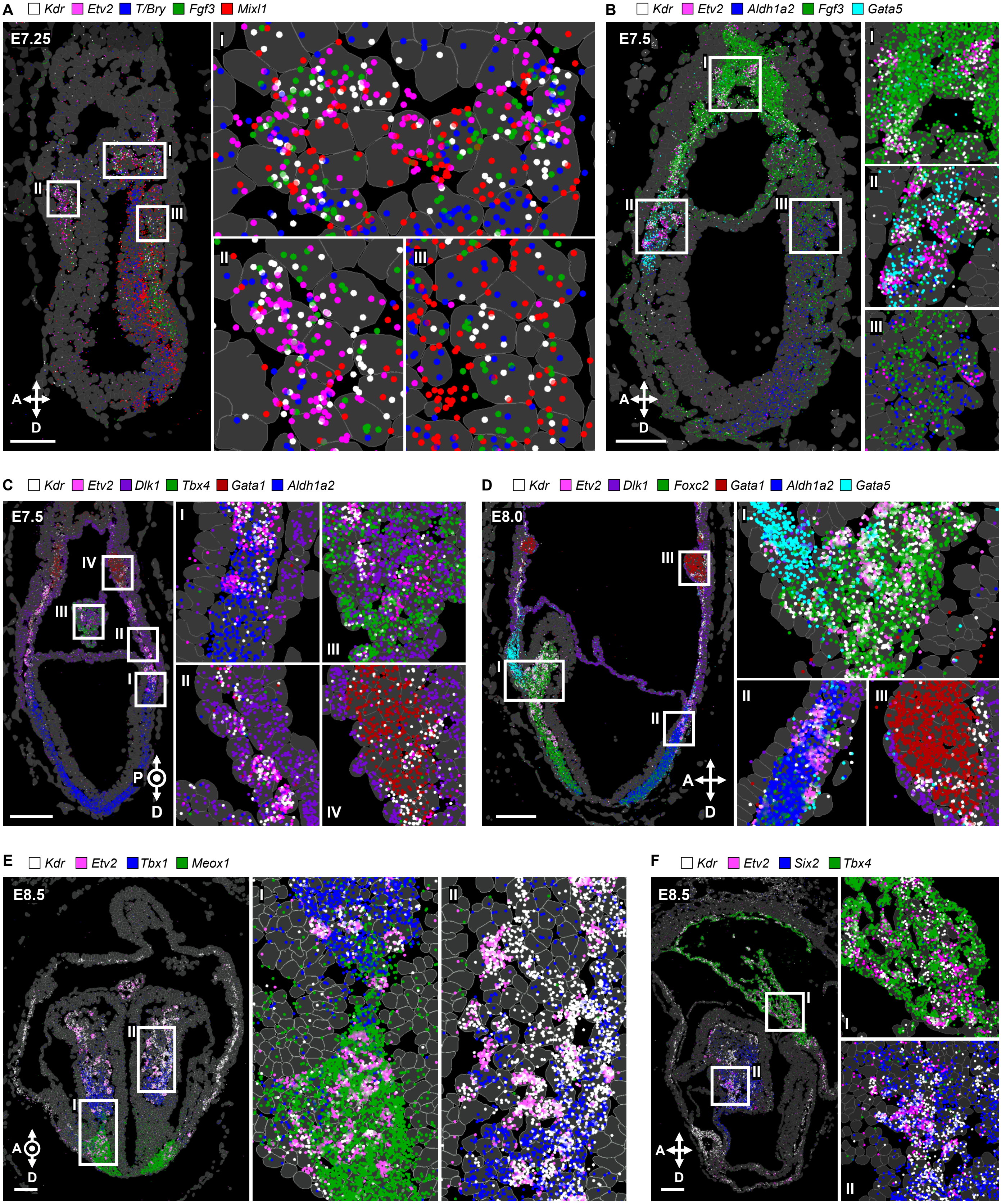
A
Kdr Etv2 T/Bry Fgf3 Mixl1
E7.25
B
Kdr Etv2 Aldh1a2 Fgf3 Gata5
E7.5
C
Kdr Etv2 Dlk1 Tbx4 Gata1 Aldh1a2
E7.5
D
Kdr Etv2 Dlk1 Foxc2 Gata1 Aldh1a2 Gata5
E8.0
E
Kdr Etv2 Tbx1 Meox1
E8.5
F
Kdr Etv2 Six2 Tbx4
E8.5

**Figure S3 | A-F,** Representative MERSCOPE sections of embryos from E7.25 to E8.5 showing *Kdr* (white) and *Etv2* (magenta) transcripts together with stage-specific local-mesoderm markers, at single-cell resolution; boxed regions are magnified on the right. Markers: **A,** E7.25: *T/Bry* (blue), *Fgf3* (green) and *Mixl1* (red). **B,** E7.5: *Aldh1a2* (blue), *Fgf3* (green) and *Gata5* (cyan). **C,** E7.5: *Dlk1* (purple), *Tbx4* (green), *Gata1* (red) and *Aldh1a2* (blue). **D,** E8.0: *Dlk1* (purple), *Foxc2* (green), *Gata1* (red), *Aldh1a2* (blue) and *Gata5* (cyan). **E,** E8.5: *Tbx1* (blue) and *Meox1* (green). **F,** E8.5: *Six2* (blue) and *Tbx4* (green). Scale bars, 100 μm. A, anterior; P, posterior; D, distal.

### Figure S4

**A** Marker genes used for correlation analysis

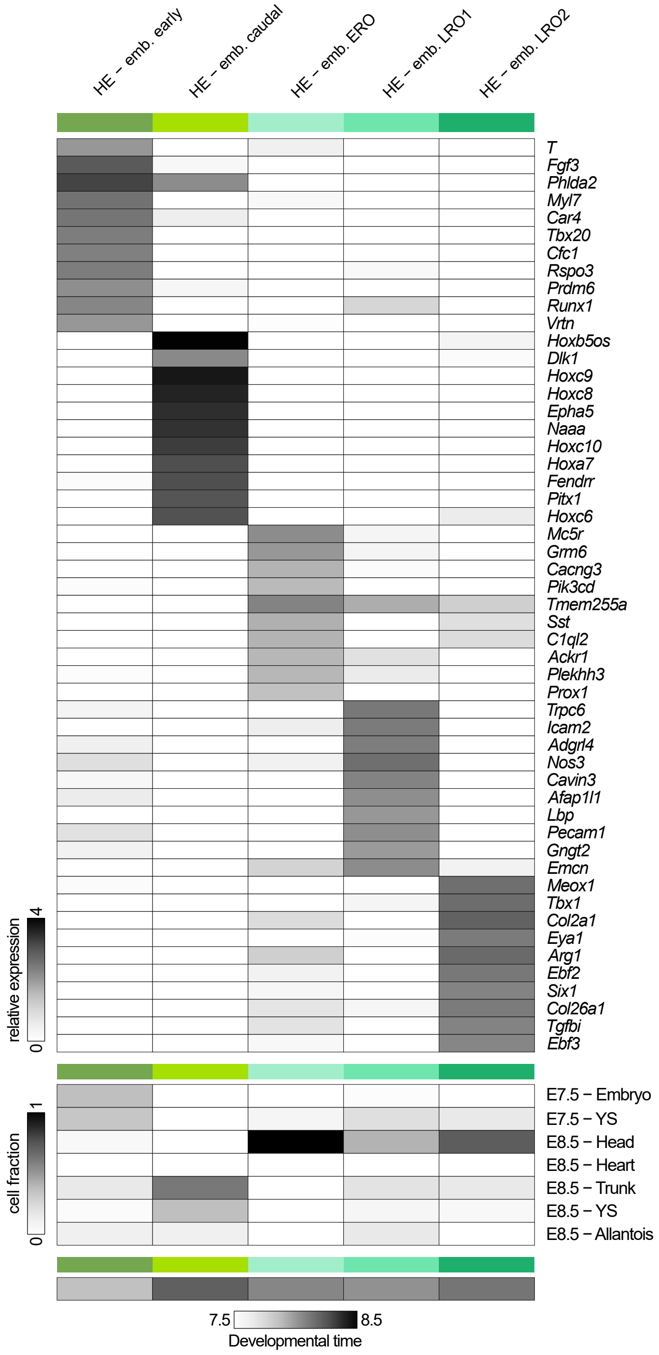


**B** Metacell expression over time

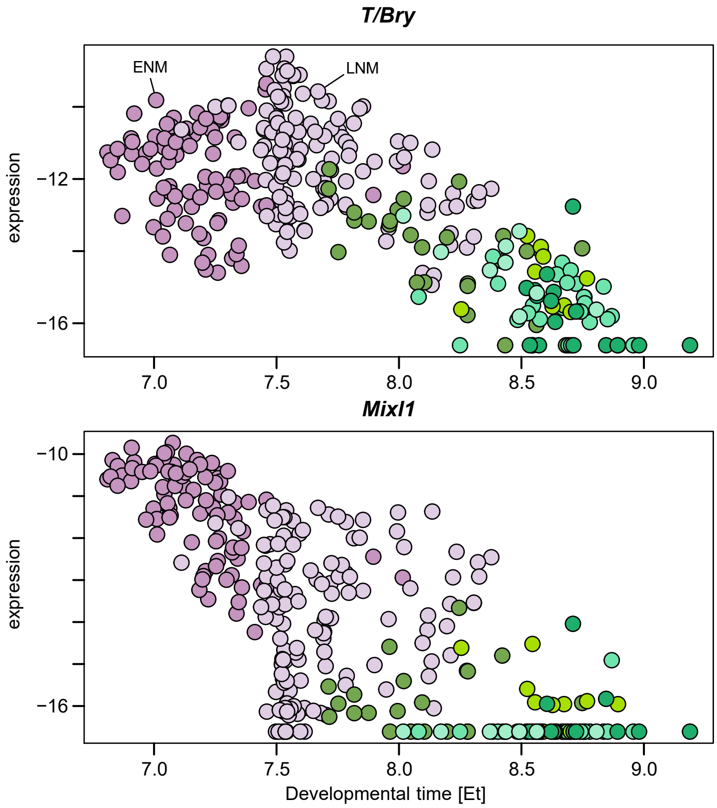

**Figure S4 | A,** Heatmap of differential marker genes showing relative expression across the embryonic HE types color coded. Bottom, the spatial distribution (cell fraction per compartment) and temporal distribution (developmental time) of each subtype. **B,** MetaCell expression of the nascent-mesoderm markers *T/Bry* and *Mixl1* across developmental time (Et), with MetaCells colored by HE subtypes and nascent mesoderm annotated in figure. HE, hematoendothelial progenitors; Emb., embryonic; ERO, early rostral; LRO, late rostral; ENM, early nascent mesoderm; LNM, late nascent mesoderm.

### Figure S5

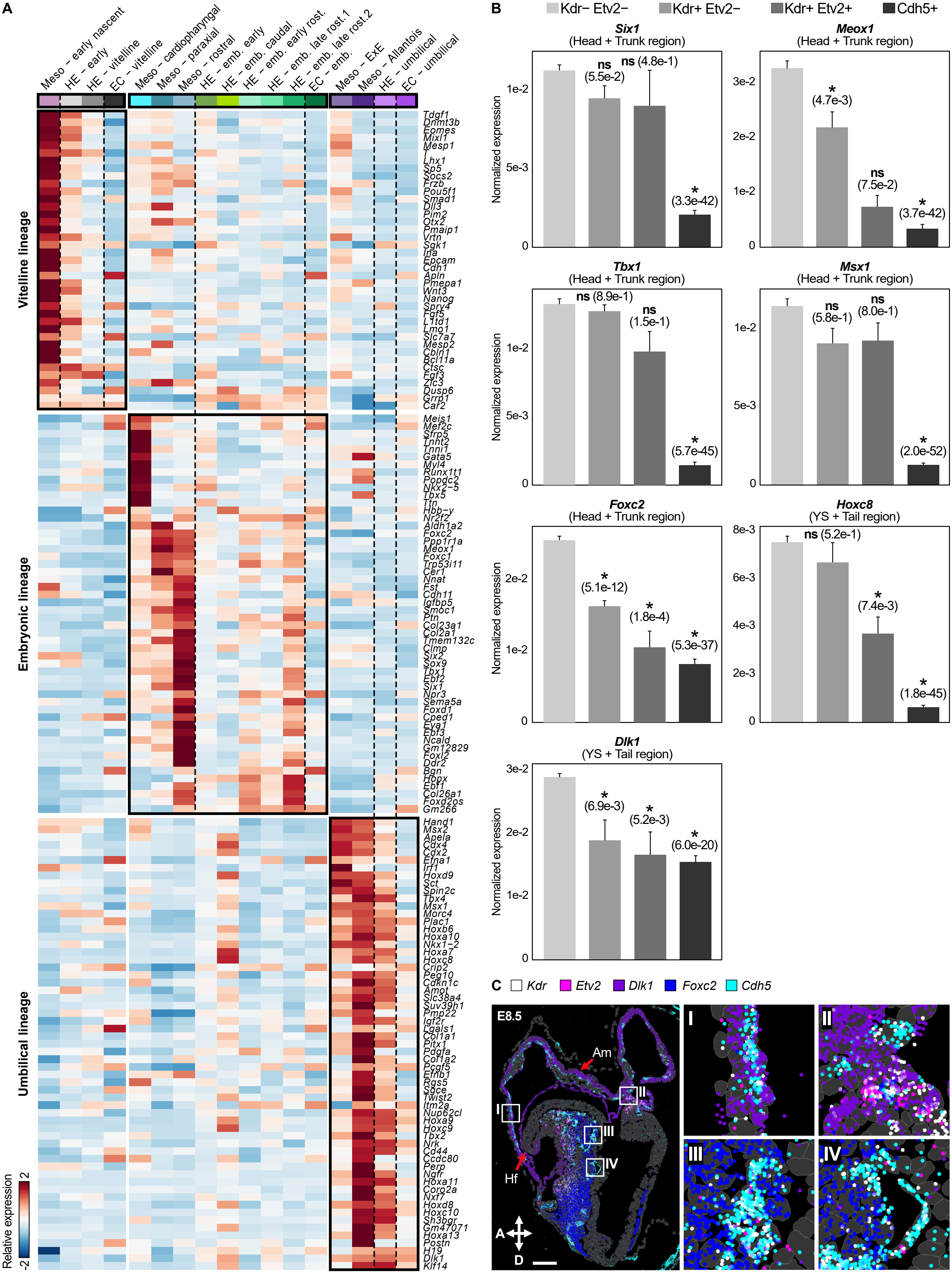
A
Meso - early nascent
HE - early
HE - vitelline
EC - vitelline
Meso - cardiopharyngeal
Meso - paraxial
Meso - rostral
HE - emb. early
HE - emb. caudal
HE - emb. early rost.
HE - emb. late rost.1
HE - emb. late rost.2
EC - emb.
Meso - ExE
Meso - Allantois
HE - umbilical
EC - umbilical
Vitelline lineage
Embryonic lineage
Umbilical lineage
Relative expression
B
Kdr− Etv2−
Kdr+ Etv2−
Kdr+ Etv2+
Cdh5+
Six1
(Head + Trunk region)
Meox1
(Head + Trunk region)
Tbx1
(Head + Trunk region)
Msx1
(Head + Trunk region)
Foxc2
(Head + Trunk region)
Hoxc8
(YS + Tail region)
Dlk1
(YS + Tail region)
Normalized expression
C
Kdr
Etv2
Dlk1
Foxc2
Cdh5
E8.5
Am
Hf
I
II
III
IV
A
D

**Figure S5 | A,** Relative expression heatmap of the differential gene-expression signatures within each of the three main vascular lineages (vitelline, embryonic and umbilical), spanning mesoderm progenitors, HEs and ECs. Boxes highlight the lineage-specific signature blocks. Separation of cell type groups indicated by dashed lines. **B,** Normalized expression of local-mesoderm markers from MERSCOPE data across populations, comparing *Kdr*-/*Etv2*-negative (local mesoderm), *Kdr*-positive/*Etv2*-negative (vascular-primed mesoderm), *Kdr*-/*Etv2*-positive (specified HE) and *Cdh5*-positive (differentiated EC) populations. Bars show the mean; adjusted p-values (Wilcoxon rank-sum test, adjusted with Benjamini-Hochberg procedure) are indicated in parentheses; asterisk, q-value < 0.05; ns, not significant. **C,** Representative MERSCOPE section of an E8.5 embryo showing *Etv2* (magenta), *Kdr* (white) and the local markers *Dlk1* (purple) and *Foxc2* (blue) within *Cdh5*-positive (cyan) ECs. Boxed regions I-IV are magnified on the right. Scale bar, 100 µm. Am, amnion; Hf, head fold; A, anterior; D, distal.

### Figure S6

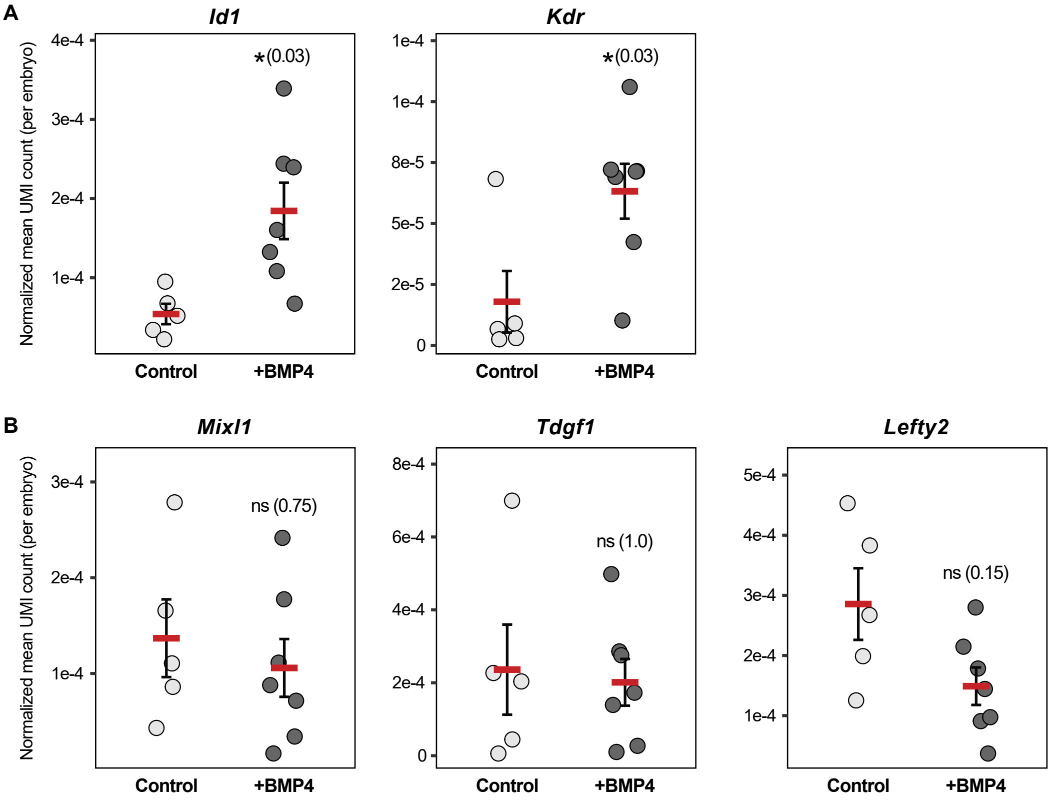
A
Id1
Kdr
Normalized mean UMI count (per embryo)
4e-4
3e-4
2e-4
1e-4
*(0.03)
1e-4
1e-4
8e-5
5e-5
2e-5
0
Control
+BMP4
B
Mixl1
Tdgf1
Lefty2
3e-4
2e-4
1e-4
ns (0.75)
8e-4
6e-4
4e-4
2e-4
0
ns (1.0)
5e-4
4e-4
3e-4
2e-4
1e-4
ns (0.15)

**Figure S6 | A-B,** Normalized mean UMI count per embryo for *Id1* and *Kdr* (**A**) and *Mixl1*, *Tdgf1* and *Lefty2* (**B**) in nascent mesoderm cells of BMP4-treated and control embryos. Each dot represents one embryo; red bars and whiskers indicate the mean and IQR, respectively. Adjusted p-values (Wilcoxon rank-sum test, adjusted with Benjamini-Hochberg procedure) are indicated in parentheses; asterisk, q-value < 0.05; ns, not significant.

### Figure S7

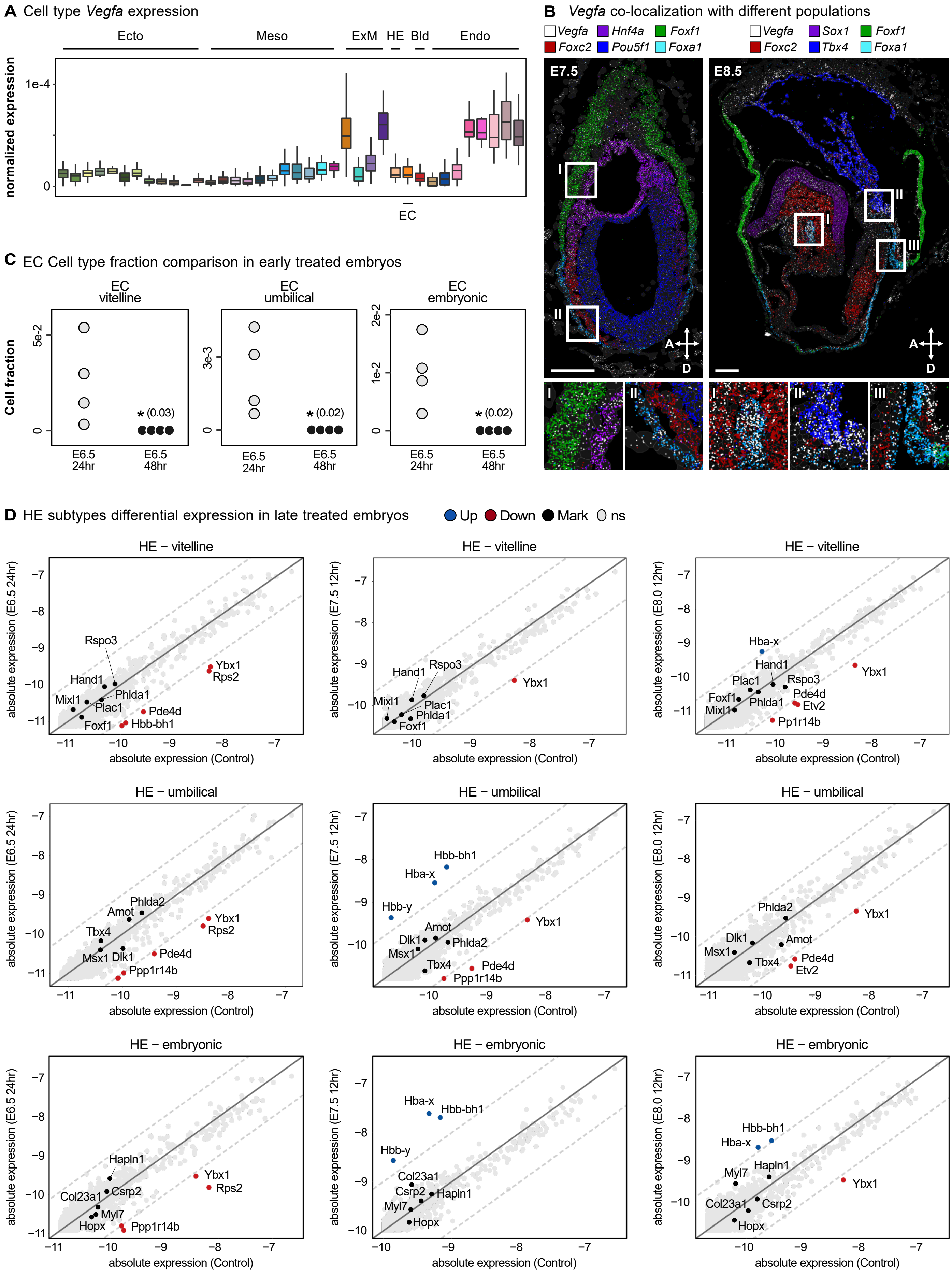

A Cell type Vegfa expression
Ecto
Meso
ExM
HE
Bld
Endo
normalized expression
1e-4
0
EC
B Vegfa co-localization with different populations
Vegfa
Hnf4a
Foxf1
Foxc2
Pou5f1
Foxa1
Sox1
Tbx4
E7.5
E8.5
I
II
III
A
D
C EC Cell type fraction comparison in early treated embryos
EC vitelline
EC umbilical
EC embryonic
Cell fraction
5e-2
3e-3
2e-2
1e-2
* (0.03)
* (0.02)
E6.5 24hr
E6.5 48hr
D HE subtypes differential expression in late treated embryos
Up
Down
Mark
ns
HE − vitelline
HE − umbilical
HE − embryonic
absolute expression (E6.5 24hr)
absolute expression (E7.5 12hr)
absolute expression (E8.0 12hr)
absolute expression (Control)
Rspo3
Hand1
Mixl1
Phlda1
Plac1
Foxf1
Ybx1
Rps2
Pde4d
Hbb-bh1
Hba-x
Etv2
Pp1r14b
Phlda2
Amot
Tbx4
Msx1
Dlk1
Ppp1r14b
Hbb-y
Hapln1
Csrp2
Col23a1
Myl7
Hopx

**Figure S7 | A,** Normalized *Vegfa* expression across cell types grouped. Boxes show the mean and IQR. **B,** Representative MERSCOPE sections showing *Vegfa* co-localization with population markers at E7.5 (left) and E8.5 (right). Channel colors are indicated in the key. Boxed regions are magnified below. Scale bars, 100 µm. A, anterior; D, distal. **C,** Fractions of vitelline, umbilical and embryonic EC in individual embryos after 24 hours (hr) or 48 hr continuous VEGFR inhibition from E6.5. Each dot represents one embryo, and adjusted p-values (Wilcoxon rank-sum test, adjusted with Benjamini-Hochberg procedure) are indicated in parentheses; asterisk, q-value < 0.05; ns, not significant. **D,** Absolute gene expression in vitelline, umbilical and embryonic HE (rows) of VEGFR-inhibited versus control embryos under three temporally restricted inhibition conditions (E6.5 24hr, E7.5 12hr and E8.0 12hr; columns), plotting treated (y axis) against control (x axis). Each point is a gene, colored by differential expression: upregulated (Up, blue), downregulated (Down, red), additionally highlighted (Mark, black) and non-significant (ns, grey). The solid line marks x = y and dashed lines a 2-fold change. Ecto, ectoderm; Meso, mesoderm; ExM, extraembryonic mesoderm; HE, hematoendothelial progenitors; Bld, primitive blood; Endo, endoderm; EC, endothelial cells.
